# START domains generate paralog-specific regulons from a single network architecture

**DOI:** 10.1101/2024.09.09.612080

**Authors:** Ashton S. Holub, Sarah G. Choudury, Ekaterina P. Andrianova, Courtney E. Dresden, Ricardo Urquidi Camacho, Igor B. Zhulin, Aman Y. Husbands

## Abstract

Functional divergence of transcription factors (TFs) has driven cellular and organismal complexity throughout evolution, but its mechanistic drivers remain poorly understood. Here we test for new mechanisms using CORONA (CNA) and PHABULOSA (PHB), two functionally diverged paralogs in the CLASS III HOMEODOMAIN LEUCINE ZIPPER (HD-ZIPIII) family of TFs. We show that virtually all genes bound by PHB (∼99%) are also bound by CNA, ruling out occupation of distinct sets of genes as a mechanism of functional divergence. Further, genes bound and regulated by both paralogs are almost always regulated in the same direction, ruling out opposite regulation of shared targets as a mechanistic driver. Functional divergence of CNA and PHB instead results from differential usage of shared binding sites, with hundreds of uniquely regulated genes emerging from a commonly bound genetic network. Regulation of a given gene by CNA or PHB is thus a function of whether a bound site is considered ‘responsive’ versus ‘non-responsive’ by each paralog. Discrimination between responsive and non-responsive sites is controlled, at least in part, by their lipid binding START domain. This suggests a model in which HD-ZIPIII TFs use information integrated by their START domain to generate paralog-specific transcriptional outcomes from a shared network architecture. Taken together, our study identifies a new mechanism of HD-ZIPIII TF paralog divergence and proposes the ubiquitously distributed START evolutionary module as a driver of functional divergence.

## INTRODUCTION

Life depends on careful control of cellular decisions. At the heart of this control is regulation of gene expression by transcription factors (TFs). Intricacy of gene regulation scales with organismal complexity^1,2,3,4,5,6^, as evidenced by the expansion in both the number and diversity of TFs during plant and animal evolution^7^. This expansion was mediated primarily by gene duplications which afforded organisms certain evolutionary advantages. For instance, paralogous TFs could retain partial or complete functional redundancy^8^. This lends robustness to biological systems, permitting organisms to better tolerate genic or environmental perturbations^9^. Alternatively, paralogous TFs could undergo functional divergence via subfunctionalization or neofunctionalization^8^. In the case of subfunctionalization, degeneration partitions the regulatory properties of a single, ancestral TF across its paralogs. During neofunctionalization, one paralog retains the ancestral function while the other takes on new roles. This functional divergence broadens the regulatory landscape of the TF family and is a central driver of cellular and organismal complexity^8,10^. Identifying mechanisms of TF functional divergence is thus key to understanding how organisms execute specific decisions ranging from stress response to immunity to development. This is a particularly challenging problem in the plant lineage, whose TF families underwent a more dramatic expansion than other kingdoms^11,12^.

One well-established mechanism of TF divergence is the occupation of different *in vivo* binding sites^1,2,3,4,5,6,13,14,15^. The acquisition of new binding sites can be influenced by several factors such as chromatin accessibility^16^, distribution of co-factors^14,17,18^ , and preferences in DNA sequence and/or shape^1,19^. Another method by which paralogs diverge is ‘antifunctionalization’, wherein paralogous TFs act antagonistically on the same pathway^20^. For instance, FLOWERING LOCUS T (FT) and TERMINAL FLOWER1 (TFL1) are paralogous TFs that promote or repress flowering, respectively^21^. Mechanistically, this is accomplished by mutually exclusive interactions between FT or TFL1 and their bZIP co-factor FD such that FD-FT complexes activate targets and FD-TFL1 complexes repress them^22^. In another example, the metazoan early gene 2 factor (E2F) family of activators and repressors similarly act antagonistically on a shared set of targets. Unlike FT and TFL1 however, their promoter binding events are temporally separated throughout the cell cycle^112^. There are likely many more biological strategies to generate paralog-specific transcriptional outputs, and plant evolutionary history suggests this kingdom is a particularly fertile ground in which to discover them^11,12^.

The CLASS III HOMEODOMAIN LEUCINE ZIPPER (HD-ZIPIII) family of plant TFs arose ∼725 million years ago and proliferated over the course of evolution. The model plant *Arabidopsis thaliana* has five paralogs *– REVOLUTA (REV), PHABULOSA (PHB), PHAVOLUTA (PHV), CORONA (CNA),* and *ATHB-8* – that collectively impact nearly all aspects of development^23,24,25,26,27,28,29,30^ (**Fig. 1a**. HD-ZIPIII paralogs are divided into two sub-clades – the REVOLUTA clade and the CORONA clade – with members displaying both functional redundancy and functional divergence. Examples of redundancy include *PHB, PHV, REV*, and *CNA* redundantly promoting dorsal identity in lateral organs and xylem identity in the vasculature^24,31,32^. An example of divergence is *CNA* antagonizing the stem cell niche, while *REV* promotes its formation or maintenance^25^. This functional divergence is driven by substitutions in the coding sequences of *REV* and *CNA*, rather than modifications at the *cis*-regulatory level, however the nature of the underlying mechanism remains unknown^25^.

**Figure 1.**
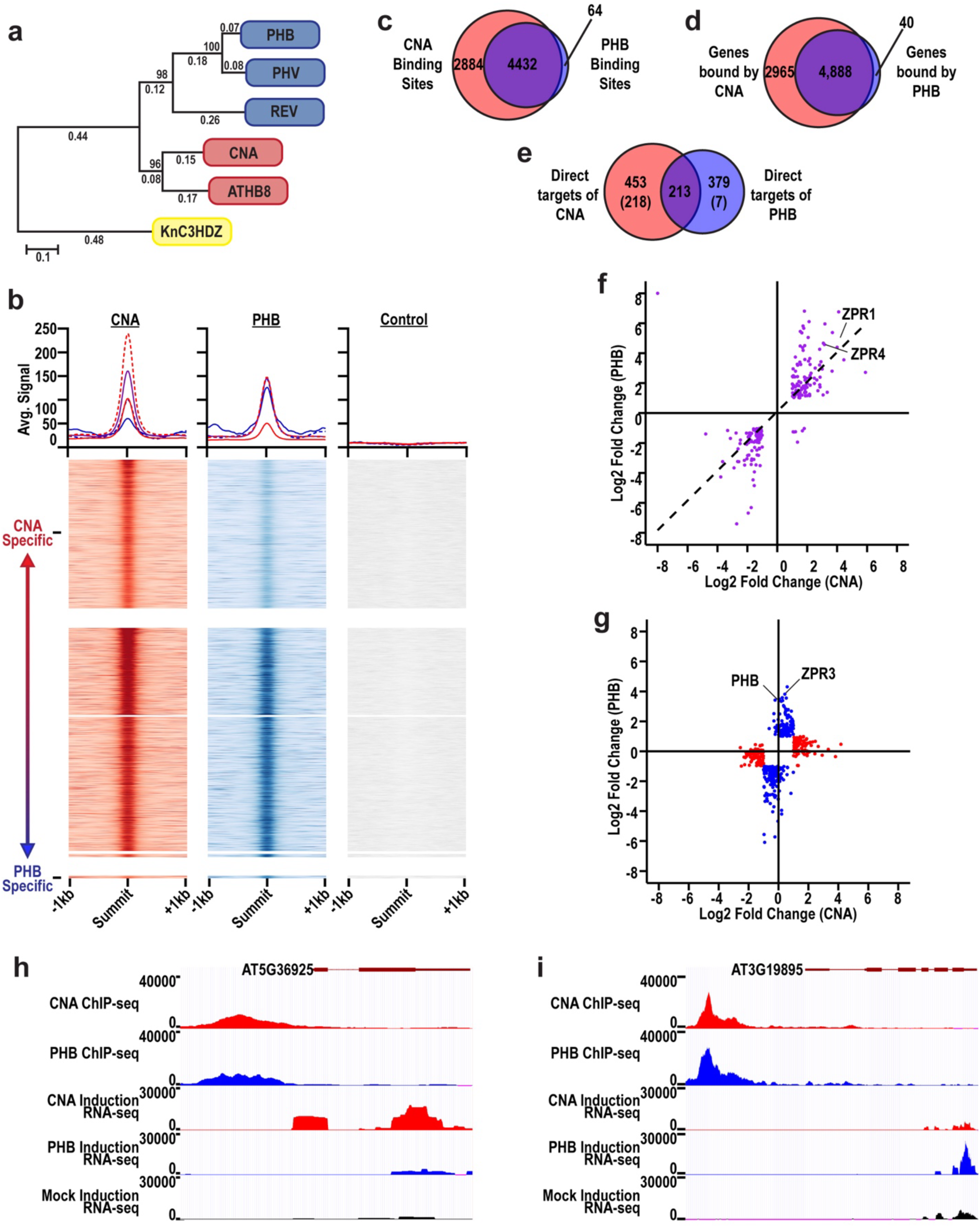
PHB and CNA have hundreds of uniquely regulated targets despite extensive overlap in genomic occupancy. **a**, Phylogenetic tree of the HD-ZIPIII family in *A. thaliana* (blue = REV subclade, red = CNA subclade). Tree rooted to the *K. nitens* HD-ZIPIII protein (KnC3HDZ, yellow). **b**, Histograms and heatmaps of ChIP-seq signal intensities compared to a non-transgenic control. Histogram lines and colors delineate five categories of binding sites: CNA specifically bound (red, solid); mutually bound – CNA higher affinity (red, dashed); mutually bound – equal affinities (purple); mutually bound – PHB higher affinity (blue, dashed), and PHB specifically bound (blue, solid). Heatmaps are separated into three major categories: CNA uniquely bound (top), mutually bound (center), and PHB uniquely bound (lower). Mutually bound heatmaps contain three further subcategories: mutually bound – CNA higher affinity (top); mutually bound – equal affinities (center); mutually bound – PHB higher affinity (lower). **c**, Venn diagram of sites bound by CNA and PHB. **d,** Venn diagram of genes bound by CNA and PHB. **e,** Venn diagram of direct targets of CNA and PHB, i.e. bound in ChIP-seq and differentially expressed in RNA-seq. Numbers in parentheses correspond to direct targets bound specifically by CNA or PHB. **f**, Scatterplot showing differential expression of genes bound and regulated by both CNA and PHB. **g**, Scatterplot showing differential expression of genes bound by both paralogs but uniquely regulated by CNA (red) or PHB (blue). **h**, **i**, Representative genome browser shots illustrating differential usage of shared binding sites: AT2G22860 is bound by both paralogs but uniquely regulated by CNA (**h**), while AT3G19895 is bound by both paralogs but uniquely regulated by PHB (**i**).

Consistent with their critical role in development, HD-ZIPIII TFs are regulated by numerous inputs including microRNAs and interacting proteins^33,34,35,36^. Additional points of regulation are suggested by their characteristic architecture, which consists of a homeodomain (HD), a leucine zipper (LZ), a MEHKLA/PAS-like domain, and a StAR-related transfer (START) domain. The latter is part of the StARkin domain superfamily, named for their kinship to steroidogenic acute regulatory protein (StAR). StARkin domains are present throughout the tree of life and are defined by a conserved α/β helix grip fold structure^37,38,39^. Dysregulation of StARkin proteins has profound effects on disease, stress responses, and development across Eukaryota^38,39,40,41,42,43,44^. StARkin domains bind a variety of lipid ligands that trigger conformational changes to modulate the activity of StARkin proteins using context-dependent regulatory mechanisms ranging from protein turnover to subcellular localization to homomeric and heteromeric complex stoichiometry (reviewed in ^45^). The diverse and varied architecture of HD-ZIPIII proteins facilitates the integration of multiple regulatory inputs and presents numerous opportunities to drive functional divergence, as amino acid substitutions underlying paralog-specific responses are not limited to DNA-binding domains^13,46,47^. HD-ZIPIII TFs are thus an excellent model to identify new regulatory mechanisms driving TF paralog divergence at the protein level.

Using genetic and genomic analyses, we find that the functional divergence of CNA and PHB is not explained by binding to different loci or opposite regulation of shared targets. Rather, the primary mechanism by which these HD-ZIPIII paralogs generate distinct transcriptional outcomes is differential usage of shared binding sites. For instance, we show that the binding profiles of CNA and PHB are nearly overlapping. Despite this, CNA and PHB each have hundreds of uniquely regulated direct targets. These unique targets occasionally include genes not bound by the other paralog; however, this is the minority case. Paralog-specific regulons are thus primarily a function of whether a given bound site is considered responsive versus non-responsive. Using deletions, chimeras, and targeted amino acid substitutions, we further demonstrate that this differential usage of shared binding sites is driven, at least in part, by the HD-ZIPIII START domain. Taken together, our study identifies a new mechanism of HD-ZIPIII TF paralog divergence and proposes the ubiquitously distributed START evolutionary module as a driver of functional divergence.

## RESULTS

### CNA and PHB bind a largely overlapping set of genomic regions

Binding new sites in the genome is a well-known mechanism by which paralogs generate distinct outcomes^1,2,3,4,5,6^. Occupation of different loci by CNA and PHB is thus a logical explanation for their functional divergence. To test this possibility, we identified genomic regions bound *in vivo* by each TF using an established short-term estradiol induction system and chromatin immunoprecipitation followed by sequencing^48^ (ChIP-seq; **Extended Data** Fig. 1). CNA and PHB bound 7,316 and 4,496 sites in the genome, corresponding to 7,853 and 4,928 genes, respectively (**Figs. 1b-d**). Remarkably, 99.2% of genes bound by PHB (4,888 out of 4,928) are also bound by CNA, with CNA uniquely occupying an additional 2,965 genes. Supporting the relevance of our findings, centrally localized partial HD-ZIPIII binding sites emerged as consensus motifs for CNA and PHB^49,50^ (VTAATNATTAB for CNA; TAATRATKATD for PHB; **Extended Data** Fig. 2). However, these motifs do not strictly predict CNA and PHB binding. For instance, VTAATNATTAB occurs 20,512 times in the *Arabidopsis* genome, yet only ∼7% of these motifs are bound by CNA (1,436 out of 20,512; **Supp. Table. 1c**). Additional elements are thus likely to refine binding site selection. A potential candidate to provide this specificity is the three-dimensional shape of DNA which plays important roles in determining *in vivo* TF binding^19,51,52^. Consistent with this prediction, DNA at bound versus unbound motifs showed clear differences in numerous shape features. For instance, comparisons of bound versus unbound motifs (and their flanking regions) uncovered strong deviations in minor groove width, helix twist, propellor twist, and roll for both CNA and PHB (**Extended Data** Fig. 3). Thus, CNA and PHB appear to bind a specific but largely overlapping set of sites, whose selection is guided by both shape and sequence of DNA.

Signal intensities at these mutually bound sites fall into three categories: higher signal for CNA (3,512 out of 9,571), higher signal for PHB (272 out of 9,571), and equivalent signal for both TFs (5,787 out of 9,571; **Fig. 1b left, middle**). CNA thus appears to bind with higher affinity than PHB at many, but not all, regions of the genome. Interestingly, PHB ChIP signal intensities are slightly higher at the summits of CNA-specific binding sites than the flanking regions (**Fig. 1b middle**). Further, signal intensities at these regions are significantly higher than those from an immunoprecipitated non-transgenic control (**Fig. 1b right; Supp. Table. 2a).** Thus, it is technically possible that PHB also binds to ‘CNA-specific’ sites, but its binding fails to meet our stringent statistical criteria. If so, CNA and PHB would occupy an essentially overlapping set of sites in the genome. Although we proceed under the assumption of CNA-specific binding, we note this alternate possible conclusion.

Our findings reveal a substantial overlap in the genomic occupancy of CNA and PHB. This observation is consistent with their partial functional redundancy^25^. Further, the unique occupancy at 2,965 additional genes by CNA provides a potential explanation for its functional divergence from PHB. By contrast, the binding profile of PHB is essentially a fully contained subset of the binding profile of CNA (**Fig. 1d**). Functional divergence of PHB activity must therefore derive from an alternate mechanism.

### CNA and PHB have hundreds of uniquely regulated direct targets despite substantial overlap in genomic occupancy

TF occupancy at a gene is not necessarily indicative of regulation (reviewed in ^53^). We therefore tested whether binding by CNA and PHB leads to changes in gene expression using short-term estradiol induction followed by transcriptome profiling^48^. After induction, 1,464 and 1,686 genes were differentially expressed in CNA- and PHB-expressing lines, respectively (**Supp. Table 3a,b**). Genes that were also bound in ChIP assays were called as direct targets. This corresponds to 666 and 592 high-confidence direct targets of CNA and PHB, respectively, which fall into three categories: CNA-specific, PHB-specific, and mutually regulated. Importantly, these datasets include known HD-ZIPIII direct targets such as the *LITTLE ZIPPER (ZPR)* genes, supporting the validity of our approach.

Of the 453 CNA-specific direct targets, 218 were from loci uniquely bound by CNA (**Fig. 1e,g**). Thus, one mechanism by which CNA generates its specific outcomes is the selection and regulation of new loci (or possibly the retention of ancestral binding sites no longer recognized by PHB). A second potential mechanism could be opposite regulation of shared direct targets^20,54^. However, this does not appear to be the case, as the 213 mutual targets of CNA and PHB are almost all regulated in the same direction at the same magnitude (**Fig. 1f**). A third potential mechanism is suggested by the fact that the remaining 240 CNA-specific direct targets are bound by both TFs but not differentially expressed in PHB-expressing lines. Similarly, PHB has 379 unique direct targets despite sharing a remarkable 99.2% of bound genes with CNA (**Figs. 1d,e,g**). This suggests differential usage of shared binding sites as a mechanism by which CNA and PHB generate specific transcriptional outcomes (illustrated in **Figs. 1h,i**). For each paralog, shared binding sites were termed ‘responsive’ if occupation promoted or inhibited transcription versus ‘non-responsive’ if occupation did not alter gene expression.

One hypothesis is that binding affinity, inferred by intensity of ChIP signal^53,55,56,57^, positively correlates with paralog-specific usage of a given binding site. To test this, we focused on two specific predictions of the hypothesis. The first states that CNA and PHB mutual direct targets should derive from genes containing shared binding sites with equivalent ChIP signal intensities (**Fig. 1d**). However, only 38% of mutual direct targets (81 out of 213) emerge from this set of genes. This set of genes also contributes 24% of CNA (107 out of 453) and 34% of PHB (129 out of 379) uniquely regulated targets, further arguing against binding affinity as a predictor of site identity (**Extended Data** Fig. 4a). A second prediction states that mutually bound, but uniquely regulated targets, should possess binding sites with ChIP signal intensities that are higher for the TF uniquely regulating the gene. We formally tested this prediction by focusing first on mutually bound genes whose shared binding sites have a higher ChIP signal for CNA (1,746 out of 4,888 bound genes). If affinity correlates with regulation, these sites should contribute most or all CNA unique targets. However, these sites contributed 12% of CNA uniquely regulated targets (54 out of 453), 27% of PHB uniquely regulated targets (101 out of 379), and 25% of mutually regulated targets (53 out of 213), arguing against this hypothesis (**Extended Data** Fig. 4b). In addition, a Spearman’s rank correlation test found no relationship between the magnitude of gene expression changes and the intensity of CNA ChIP signals (ρ = 0.24, p = 1; **Extended Data** Fig. 5a). Similar analyses found that PHB ChIP signals also failed to correlate with magnitude of gene expression changes (ρ = 0.26, p = 0.99; **Extended Data** Fig. 5b). These analyses suggest binding affinity does not dictate responsive versus non-responsive site identity. To test this further, we used additional orthogonal tests to determine whether ChIP signals at responsive sites (i.e. in direct targets) are higher than those at non-responsive sites (i.e. in non-regulated genes). The empirical distribution of ChIP signals deviated for both CNA and PHB (Kruskal-Wallis p-values: 0.006 (CNA), 0.007 (PHB); **Extended Data** Fig. 6). However, differences in mean ChIP signal were small, showing a slight increase only in upregulated versus non-regulated genes (Mann-Whitney *q*-values: 0.007 (CNA), 0.031 (PHB); **Extended Data** Fig. 6). Taken together, these qualitative and quantitative analyses suggest that paralog-specific interpretation of shared binding sites is not driven primarily by their relative binding affinities.

Supporting the biological relevance of these findings, GO term analyses of CNA-specific, PHB-specific, and mutual direct targets revealed both similarities and differences in biological processes. For instance, the most overrepresented processes regulated specifically by CNA almost all relate to wax and fatty acid biosynthesis, as well as stress responses (**Extended Data** Fig. 7). By contrast, PHB specifically downregulates several players in the auxin and brassinosteroid hormone signaling pathways (**Extended Data** Fig. 7). Processes regulated by both paralogs include photosynthesis and light responses, as well as camalexin and phytoalexin biosynthesis (**Extended Data** Fig. 7). Taken together, our analyses suggest functional divergence of HD-ZIPIII paralogs is driven primarily by differential usage of shared binding sites which appears to be independent of binding affinity.

### The START domain is required for full CNA function

Our analyses propose paralog-specific interpretation of shared binding sites contributes to the functional divergence of HD-ZIPIII TFs. Further, coding sequence swaps between HD-ZIPIII subclade members indicate their functional divergence is due in part to amino acid substitutions; the identity and position of these substitutions are unknown^25^. Notably, HD-ZIPIII TFs contain a START domain (**Fig. 2a**), known to control transcriptional regulators in plants and animals through diverse regulatory mechanisms^45,48,58,59,60,61,62^. In addition, the START domain of PHB enhances its transcriptional potency at *ZPR3* and *ZPR4* targets^48^. Thus, the START domain is an attractive candidate to drive HD-ZIPIII functional divergence through paralog-specific interpretation of shared binding sites.

**Figure 2.**
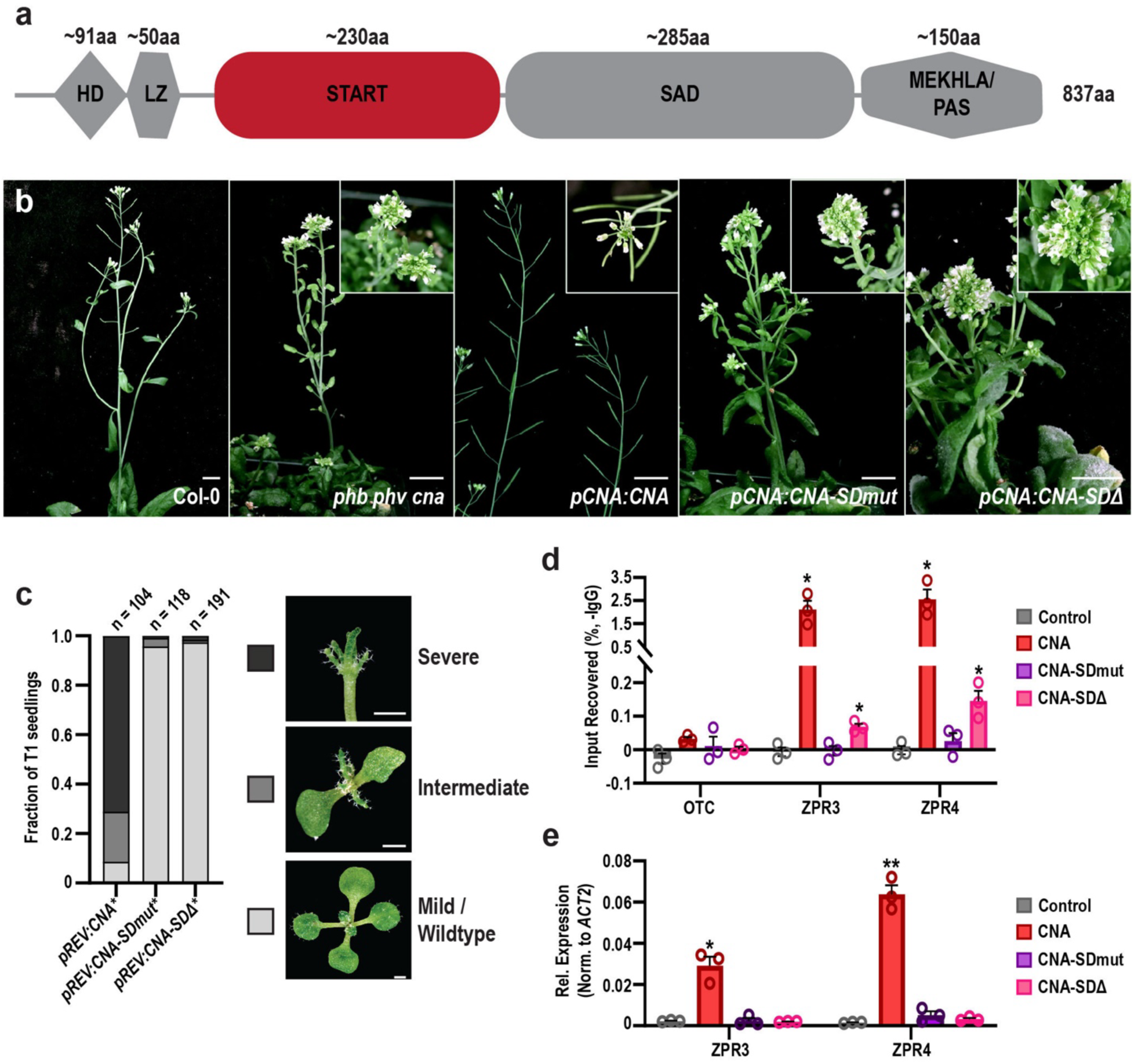
The START domain is required for the developmental activity of CNA. **a**, General structure of HD-ZIPIII proteins (HD = homeodomain, LZ = leucine zipper, START = StAR-related lipid transfer, SAD = START adjacent domain, MEKHLA/PAS). **b**, The *phb phb cna* triple mutant has a strong pleiotropic phenotype (Prigge et al., 2005). The *pCNA:CNA* transgene fully complements the mutant phenotype as plants appear wildtype, whereas the *pCNA:CNA-SDmut* and *pCNA:CNA-SDΔ* transgenes do not. **c**, Phenotypic scoring of primary transformants (*n* above bar) expressing miR-166 insensitive (*) *CNA-SDmut* and *CNA-SDΔ* under the *REV* promoter. Ectopic *CNA\** expression leads to severe (black) or intermediate (dark grey) gain-of-function phenotypes, whereas plants mis-expressing *CNA-SDmut\** or *CNA-SDΔ*\* appear wildtype (light grey). **d**, CNA and CNA-SDΔ bind to regulatory regions of ZPR3 and ZPR4 and are significantly enriched above the *ORNITHINE TRANSCARBAMYLASE (OTC)* negative control locus; note CNA-SDΔ signal is significantly lower than that of CNA. CNA-SDmut does not bind to *ZPR3* or *ZPR4* regulatory regions. **e**, CNA robustly activates *ZPR3* and *ZPR4* targets in 24 h estradiol-induction experiments, whereas CNA-SDmut and CNA-SDΔ do not. Non-transgenic control (grey), *pOlexA:CNA\** (red), *pOlexA:CNA-SDmut\** (purple), and *pOlexA:CNA-SDΔ\** (pink). Statistics are against the non-transgenic control. * = p-value ≤ 0.05, ** = p-value ≤ 0.01, Student’s t-test.

A regulatory role for the PHB START domain has been established^48^. We therefore tested whether the START domain of CNA is similarly required for its activity. To assess for START regulatory roles in development, we employed two genetic assays previously used to characterize the PHB START domain^48^. Like *PHB*, loss of function single mutants of *CNA* appear wildtype^25^. Thus, the first genetic assay involves complementing a *phb phv cna* loss-of-function mutant which has a strong pleiotropic phenotype^25^. We began by replacing the START domain of the functional *pCNA:CNA* reporter^63^ with the 21-nt microRNA166 (miR166) recognition site found within the START domain coding sequence^28^ (*pCNA:CNA-SDΔ* ). Impacts of START domain deletion can thus be assayed without confounding effects from ectopic CNA transcript accumulation^64^. In a separate construct, we introduced a set of mutations in the ligand-binding pocket that do not perturb HD-ZIPIII START secondary structure^48^ (*pCNA:CNA-SDmut*). Unlike *pCNA:CNA*, neither *pCNA:CNA-SDmut* nor *pCNA:CNA-SDΔ* rescued the *phb phv cna* mutant phenotype (**Fig. 2b**). In an orthologous genetic approach, we repurposed the established *pREV:CNA* reporter into a gain-of-function, highly sensitive readout of CNA activity by introducing a silent mutation into its miR166 recognition site^25^ (*pREV:CNA\**). As expected, over 90% of *pREV:CNA\** primary transformants show phenotypes characteristic of ectopic HD-ZIPIII activity including dorsalized leaves^27,28,64^ (**Fig. 2c**). By contrast, *pREV:CNA*-SDmut* and *pREV:CNA*-Delta* primary transformants are indistinguishable from wildtype plants (**Fig. 2c**). Our genetic assays demonstrate the START domain is required for CNA to fulfil its developmental function.

The START domain of PHB potentiates transcriptional activity using multiple distinct mechanisms prompting us to test whether the CNA START domain behaves similarly^48^. First, we tested whether the START domain impacts the ability of CNA to bind to DNA. ChIP qPCR showed strong enrichment of CNA at two *ZPR* loci (**Fig. 2d**). This binding was not observed with CNA-SDmut (**Fig. 2d**), matching the behavior of PHB variants with analogous mutations^48^. Surprisingly, CNA-SDΔ also showed strongly reduced occupancy at *ZPR3* and *ZPR4* (**Fig. 2d**). This behavior contrasts with PHB-SDΔ whose occupancy at these loci was nearly identical to PHB^48^. Thus, the impact of the START domain on DNA binding appears to differ between HD-ZIPIII paralogs. Next, we tested whether the CNA START domain affects the regulation of these targets using short-term estradiol inductions and RT-qPCR. As expected, transcript levels of *ZPR* targets were upregulated by CNA and unchanged after induction of CNA-SDmut (**Fig. 2e**). CNA-SDΔ also failed to upregulate these targets suggesting its sharp reduction in genomic occupancy at these loci has functional consequences (**Fig. 2e**). Taken together, these analyses demonstrate a role for the START domain in regulating CNA activity and suggest START-mediated effects on DNA binding differ across the HD-ZIPIII subclades.

### CNA and PHB START domains promote DNA binding and may help distinguish responsive from non-responsive binding sites

Having established a regulatory role for START domains of both paralogs at established *ZPR* targets, we next tested how their loss affects binding and regulation of CNA and PHB targets genome wide. ChIP-seq identified 3,276 and 4,490 sites bound by CNA-SDΔ and PHB-SDΔ, corresponding to 4,003 and 4,300 bound genes, respectively (**Figs. 3a-c, g-i**). These binding profiles are essentially fully contained subsets of their respective wildtype counterparts. For instance, 99.2% of genes bound by CNA-SDΔ (3,971 out of 4,003) are also bound by CNA, with CNA recognizing an additional 3,706 genes (**Fig. 3c**). In the case of PHB-SDΔ, 99.0% of bound genes (4,260 out of 4,300) are also bound by PHB, with PHB recognizing an additional 623 genes (**Fig. 3i**). Deletion of the CNA and PHB START domains thus leads to reduced genomic occupancy of these TFs, and this effect is more dramatic for CNA than PHB (**Figs. 3b,h**). Signal intensities at mutually bound sites fall into three categories: higher signal for the wildtype protein, higher signal for the START-deleted variant, and equivalent signal for both variants (**Figs. 3a,g**). Interestingly, nearly all binding sites fall in the first and last categories. This indicates occupancy at a given locus is either unaffected or reduced by START domain deletion, with reductions occurring much more often with CNA-SDΔ than PHB-SDΔ (**Figs. 3a,g**). Thus, one function of the START domain may be to increase HD-ZIPIII binding affinities in a paralog- and site-specific manner.

**Figure 3.**
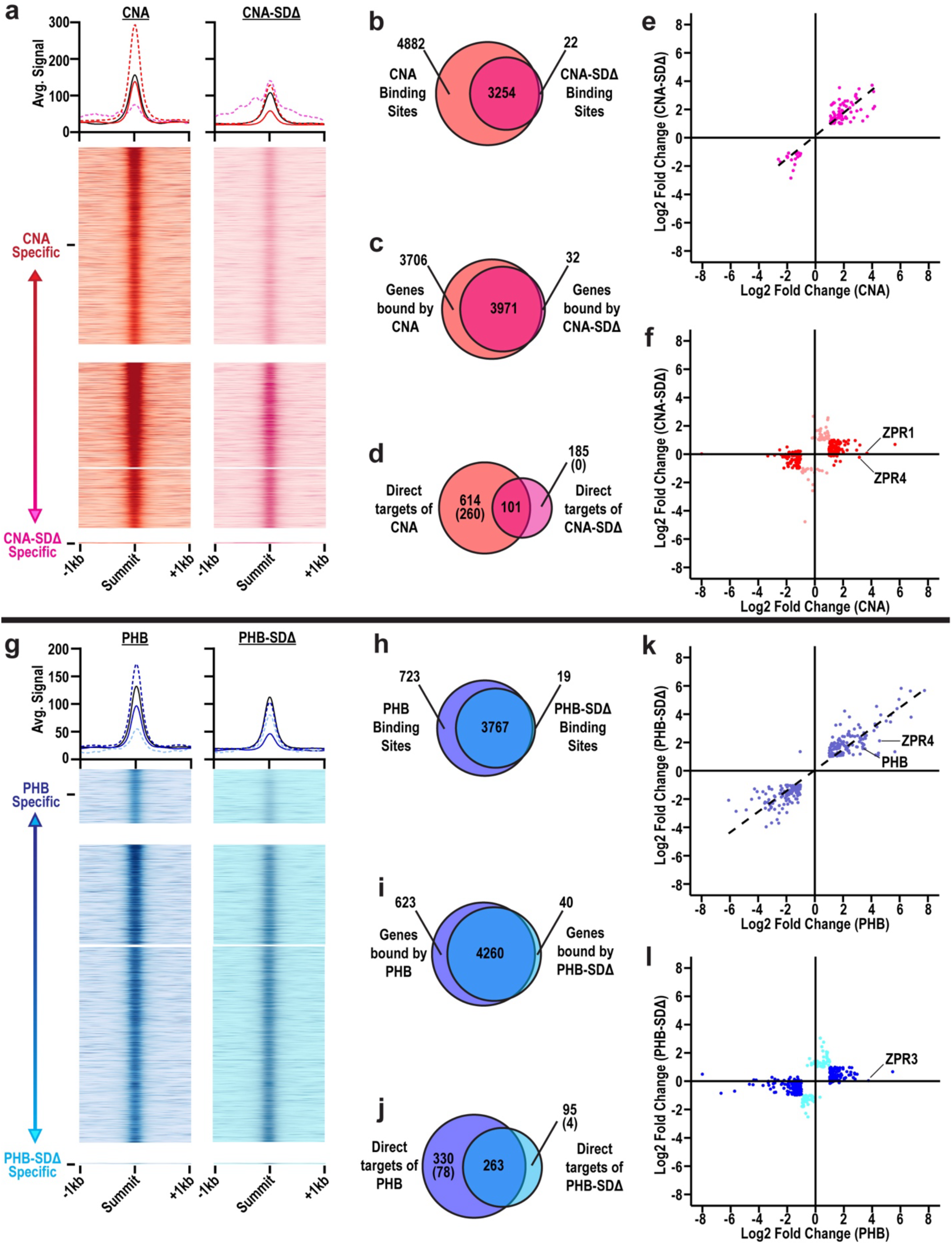
PHB and CNA START domains may help distinguish responsive from non-responsive shared binding sites. **(a-f)** CNA vs CNA-SDΔ analyses (red vs pink). (**g-l**) PHB vs PHB-SDΔ analyses (blue vs cyan). **a,g**, Histograms and heatmaps of ChIP-seq signal intensities. Lines on histograms delineate categories as described in Fig. 1. Heatmaps are separated into categories and subcategories as described in Fig. 1. **b,h,** Venn diagrams of sites bound by wildtype and START-deleted variants. **c,i,** Venn diagrams of genes bound by wildtype and START-deleted variants. **d,j,** Venn diagrams of direct targets of wildtype and START-deleted variants, i.e. bound in ChIP-seq and differentially expressed in RNA-seq. Numbers in parentheses correspond to direct targets bound specifically by wildtype or START-deleted variants. **e,k**, Scatterplots showing differential expression of genes bound and regulated by both wildtype and START-deleted variants. **f,** Scatterplot showing differential expression of genes bound by both variants but uniquely regulated by CNA (red) or CNA-Delta (pink). **l**, Scatterplot showing differential expression of genes bound by both variants but uniquely regulated by PHB (blue) or PHB-SDΔ (cyan).

If shared binding site usage is controlled by HD-ZIPIII START domains, this should be reflected in gene expression changes at commonly bound genes. To begin to assess this, we tested how loss of their START domains impacts regulation of CNA and PHB targets. After induction, 642 and 1005 genes were differentially expressed in CNA-SDΔ- and PHB-SDΔ-expressing lines, respectively (**Supp. Table 3c,d**). This corresponds to 185 and 95 high-confidence direct targets of CNA-SDΔ and PHB-SDΔ, respectively (**Figs. 3d,j**). Comparing CNA-SDΔ and PHB-SDΔ to their wildtype counterparts generates three categories of direct targets: unique to the wildtype protein, unique to the START-deleted variant, and mutually regulated (**Figs. 3d,j**). We first asked whether the START domain controls the direction of target regulation and found that mutual targets of wildtype and START-deleted variants are almost all regulated in the same direction (**Figs. 3e,k**). Interestingly, the magnitudes of gene expression changes are often lower in START-deleted samples which could be due to reduced transcriptional potency of these variants^48^. Alternatively, this may be a downstream consequence of their reduced genomic occupancy at these shared sites (**Figs. 3a,g**). We next asked whether direct targets specific to each wildtype protein are explained by their expanded genomic occupancy. Supporting this idea, 42.3% of CNA-specific targets are not bound by CNA-SDΔ (260 out of 614), and 23.6% of PHB-specific targets are not bound by PHB-SDΔ (78 out of 330). However, this is the minority case, with 57.7% of CNA-specific targets (354 out of 614) and 76.4% of PHB-specific targets (252 out of 330) also bound by their START-deleted variant (**Figs. 3d,f,j,l; Extended Data** Fig. 8). In addition, despite the binding profiles of CNA-SDΔ and PHB-SDΔ being essentially fully contained subsets of their wildtype counterparts, these proteins have 185 and 95 unique direct targets, respectively (**Figs. 3d,f,j,l; Extended Data** Fig. 8). Thus, the START domain may help CNA and PHB discriminate between responsive and non-responsive binding sites. Taken together, our transcriptional genomic assays support a role for START domains as potential drivers of HD-ZIPIII functional divergence.

### Chimeric proteins have different effects on CNA developmental activity

Deletion analyses suggest the START domain may contribute to both the selection and regulation of HD-ZIPIII targets (**Fig. 3**). One approach to generate additional support for this hypothesis is to leverage evolutionary history. HD-ZIPIII TFs are deeply conserved but contain numerous lineage specific substitutions within their START domains. This pool of natural variation can be functionally interrogated by replacing the START domain of an *Arabidopsis* HD-ZIPIII protein with START domains from orthologs in other species. The resulting chimeras probe START-directed regulatory properties while minimizing potential confounding effects associated with full domain deletions.

To guide the selection of this *Arabidopsis* HD-ZIPIII protein, we reconstructed the evolutionary history of the HD-ZIPIII family and discovered that the ancestral-most HD-ZIPIII protein most closely resembles CNA (**Extended Data** Fig. 9). CNA was thus selected as the transgenic platform for our chimeras. Orthologous START domain sequences were then chosen using horizontal and vertical approaches (**Extended Data** Figs. 9**, 10**). In the horizontal approach, the CNA START domain was replaced by START domains from orthologs in five extant species: *Zea mays* (ZmHOX29), *Selaginella moellendorffii* (SmHOX32), *Physcomitrella patens* (PpHOX32)*, Klebsormidium nitens* (KnC3HDZ), and *Chlorokybus atmophyticus* (CaC3HDZ). In the vertical approach, the CNA START domain was replaced by START domains from ancestral reconstructions of HD-ZIPIII sequences at the base of the Angiosperm (AngioAnc) or Charophyte (CharoAnc) lineages.

To assess the effect of each START domain on CNA regulation, we first modified the *pREV:CNA\** transgene from **Fig. 2c** to express miR166-resistant versions of chimeras. Using the frequency of HD-ZIPIII gain-of-function phenotypes as a readout, we found five sequences that can substitute for CNA START developmental function, and three that cannot (**Fig. 4a**). Our chimeric constructs are thus likely to be useful tools to identify START-directed effects on HD-ZIPIII divergence. Chimeras were then cloned into the estradiol-inducible system to test for effects on DNA binding and target activation. As expected, the five chimeras that condition developmental phenotypes also activate *ZPR3* and *ZPR4* targets (**Fig 4b**). Surprisingly, the three chimeras that could not affect development fell into two sub-classes. Members of one sub-class showed no activation of *ZPR3* or *ZPR4* (CNA-SmHOX32 and CNA-CaC3HDZ) while the other (CNA-KnC3HDZ) robustly activated both targets (**Fig. 4b**). We therefore selected three chimeras for further analyses: 1) CNA-SmHOX32, which does not appear to function in molecular or phenotypic assays, 2) CNA-PpHOX32, which appears to fully function in both assays, and 3) CNA-KnC3HDZ, which binds and activates *ZPR* targets yet fails to condition developmental phenotypes.

**Figure 4.**
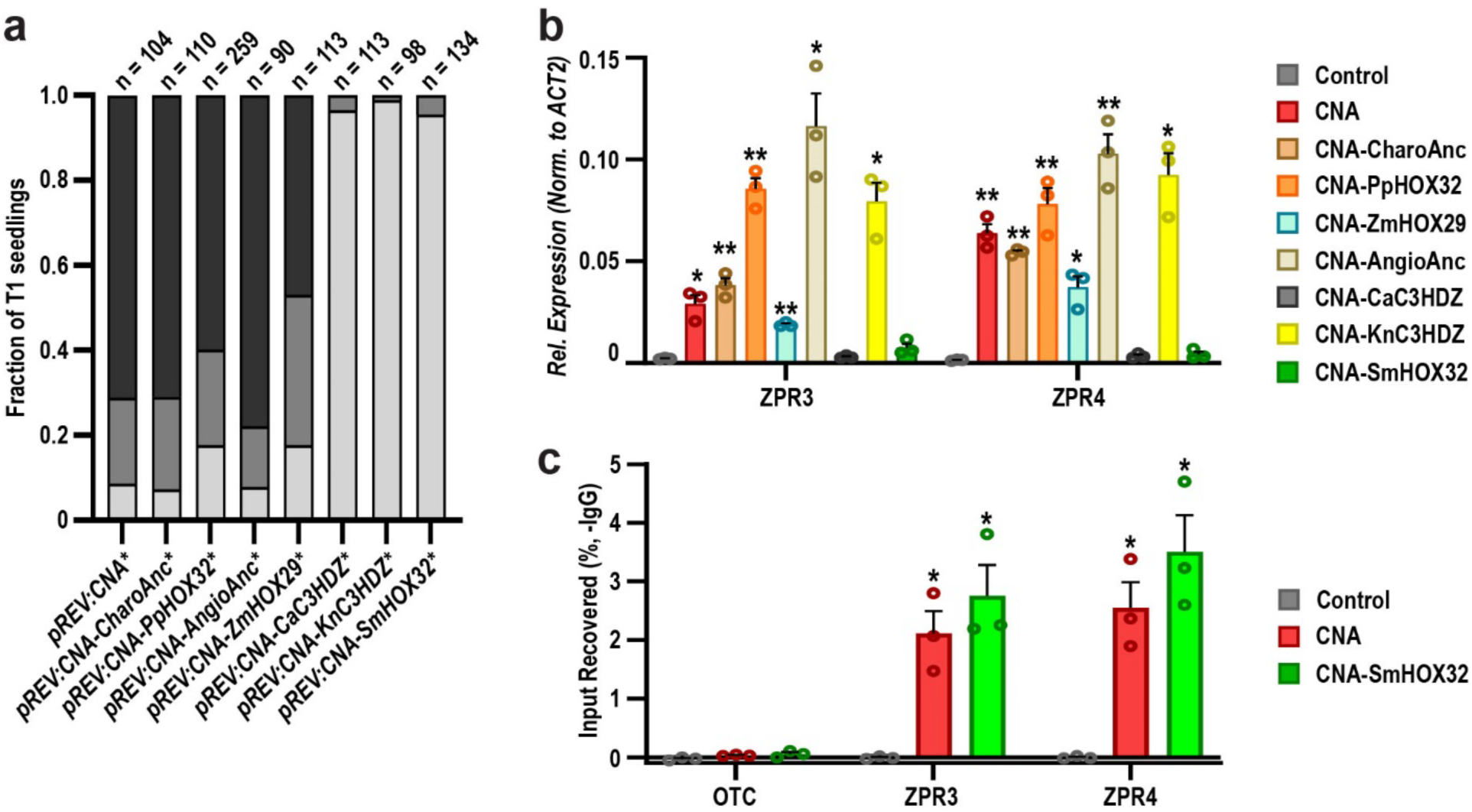
Chimeric CNA proteins have distinct effects on development and gene regulation. **a**, Phenotypic scoring of primary transformants (*n* above bar) expressing miR-166 insensitive (*) chimeric variants of *CNA* driven by the *REV* promoter. Ectopic *CNA\** expression leads to severe (black) or intermediate (dark grey) gain-of-function phenotypes, and five chimeric constructs produce similar effects (leftmost five bars). The remaining three chimeric constructs do not obviously perturb development as plants appear wildtype (light grey; rightmost three bars). Phenotypes resemble those shown in Fig. 2. **b**, CNA robustly activates *ZPR3* and *ZPR4* in 24 h estradiol-induction experiments while chimeric variants of CNA show a complex pattern of target induction. **c**, CNA and CNA-SmHOX32 both bind to regulatory regions of ZPR3 and ZPR4 and are significantly enriched above the *OTC* negative control. Nomenclature: non-transgenic control (Control), Charophyte ancestrally reconstructed variant (CharoAnc), *Physcomitrella patens* (PpHOX32), Angiosperm ancestrally reconstructed variant (AngioAnc), *Zea mays* (ZmHOX29), *Chlorokybus atmophyticus* (CaC3HDZ), *Klebsormidium nitens* (KnC3HDZ), and *Selaginella moellendorffii* (SmHOX32). * = p-value ≤ 0.05, ** = p-value ≤ 0.01, Student’s t-test.

### Chimeric proteins show differences in target selection and regulation

A simple explanation for the failure of CNA-SmHOX32 to activate *ZPR3* and *ZPR4* targets is loss of DNA binding. However, CNA-SmHOX32 ChIP qPCR enrichment values are indistinguishable from unmodified CNA at these loci (**Fig. 4c; Extended Data** Fig. 11). One possibility is this chimeric protein is non-functional and has lost the ability to regulate bound targets. Alternatively, replacing the CNA START domain with the SmHOX32 START domain may have shifted its binding profile and/or repertoire of responsive versus non-responsive sites. The latter would be consistent with results from START deletion assays (**Fig. 3**). To differentiate between these possibilities, we first identified the genomic regions bound by CNA-SmHOX32 using ChIP-seq. CNA-SmHOX32 bound 9,679 sites in the genome corresponding to 9,816 genes (**Figs. 5a-c**). Over 99.9% of these bound genes (9,809 out of 9,816) are shared with CNA, with CNA uniquely occupying an additional 2 genes (**Fig. 5c**). Thus, replacing the CNA START domain with the SmHOX32 START domain has a negligible effect on genome occupancy as CNA and CNA-SmHOX32 have virtually identical binding profiles.

**Figure 5.**
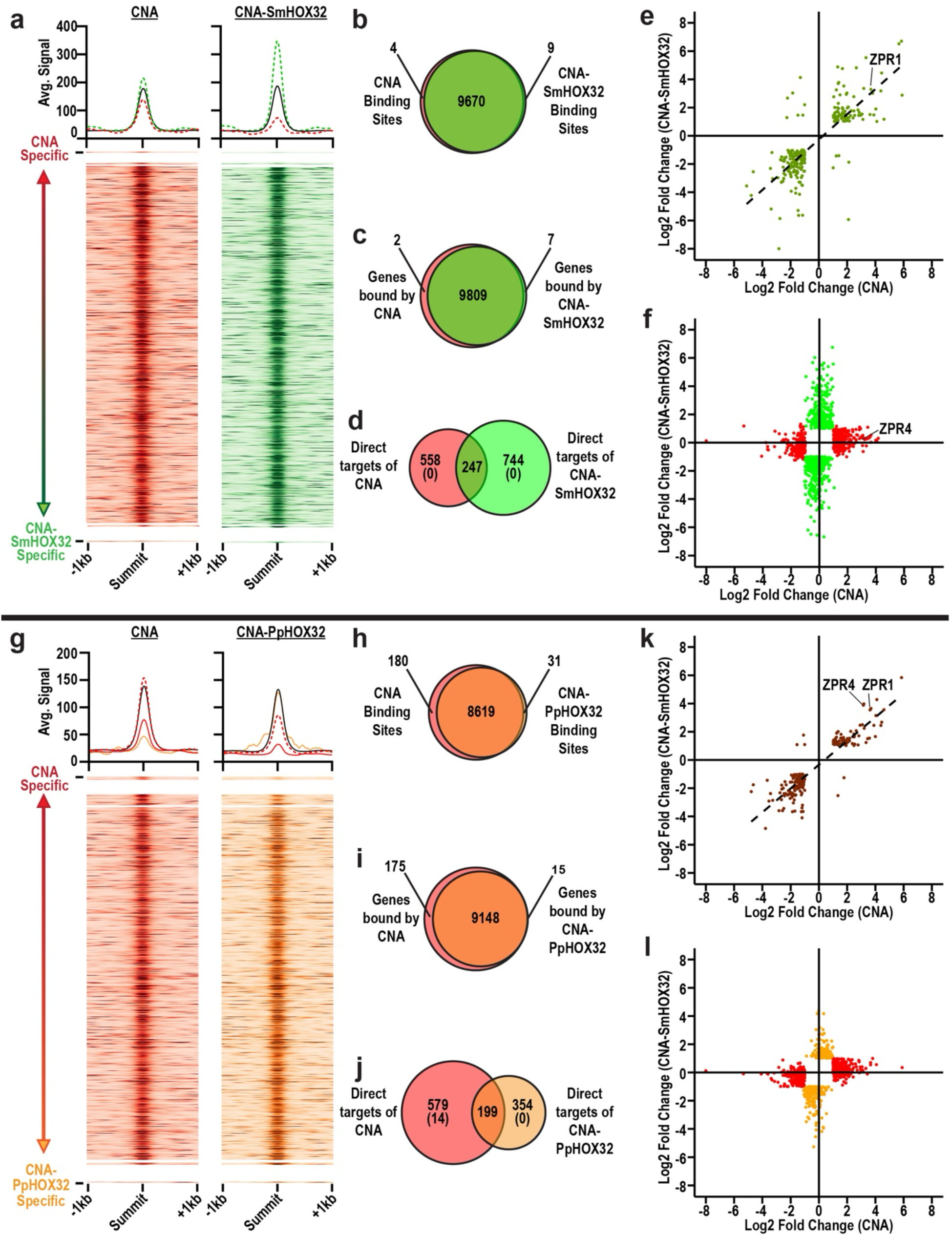
Chimeric CNA proteins show widespread differential usage of shared binding sites. (a-f) CNA vs CNA-SmHOX32 analyses (red vs green). (**g-l**) CNA vs CNA-PpHOX32 analyses (red vs orange). **a,g**, Histograms and heatmaps of ChIP-seq signal intensities. Lines on histograms delineate categories as described in Fig. 1. Heatmaps are separated into categories and subcategories as described in Fig. 1. **b,h,** Venn diagrams of sites bound by CNA and its chimeras. **c,i,** Venn diagrams of genes bound by CNA and its chimeras. **d,j,** Venn diagrams of direct targets of CNA and its chimeras, i.e. bound in ChIP-seq and differentially expressed in RNA-seq. Numbers in parentheses correspond to direct targets bound specifically by CNA or its chimeras. **e,k**, Scatterplots showing differential expression of genes bound and regulated by both CNA and its chimeras. **f,** Scatterplot showing differential expression of genes bound by CNA but uniquely regulated by CNA (red) or CNA-SmHOX32 (green). **l**, Scatterplot showing differential expression of genes bound by both variants but uniquely regulated by CNA (red) or CNA-PpHOX32 (orange).

We next tested the impact of CNA-SmHOX32 on gene expression via transcriptome profiling of CNA-SmHOX32-induced lines. After induction, 2203 genes were differentially expressed in CNA-SmHOX32-expressing lines corresponding to 991 high-confidence direct targets (**Fig. 5d; Supp. Table 3e**). CNA and CNA-SmHOX32 direct targets fall into three categories: CNA-specific, CNA-SmHOX32-specific, and mutually regulated (**Fig. 5d**). Despite binding to nearly identical sets of genes, CNA and CNA-SmHOX32 have remarkably few mutually regulated targets (**Figs. 5d-e**), with most genes showing specific regulation by either CNA (558 out of 805) or CNA-SmHOX32 (744 out of 991; **Figs. 5d,f; Extended Data** Fig. 8). Thus, replacing the CNA START domain with the SmHOX32 START domain profoundly affects target regulation, primarily through differential usage of shared binding sites.

Similar analyses found the CNA-PpHOX32 chimera bound 8,650 sites in the genome corresponding to 9,163 genes (**Figs. 5g-i**). Over 99.8% of these bound genes (9,148 out of 9,163) are shared with CNA, with CNA uniquely occupying an additional 175 genes. Pairing ChIP-seq with transcriptome profiling again found widespread differential usage of shared binding sites. For instance, despite their extensive binding overlap, CNA-PpHOX32 has 354 direct targets that are not shared with CNA (**Figs. 5j,l; Extended Data** Fig. 8). Similarly, CNA has 579 unique targets, 97.5% of which derive from mutually bound genes (565 out of 579; **Fig. 5j; Extended Data** Fig. 8).

Taken together, these developmental and functional genomics assays support the idea that START domains modulate HD-ZIPIII TF activity primarily by distinguishing whether a site is considered responsive or non-responsive by a given paralog.

### START point mutations mimicking HD-ZIPIII evolution drive differences in target selection and regulation

Domain swaps are consistent with START domains driving changes in HD-ZIPIII transcriptional outputs. However, TF evolution rarely involves wholesale exchanges of domains, instead working largely through gradual accrual of individual amino acid substitutions. If START domains are indeed drivers of HD-ZIPIII functional divergence, even minimal changes in their sequences should impact target selection and/or differential usage of shared binding sites. The CNA-KnC3HDZ chimera provides an opportunity to test this notion, as this protein retains some molecular properties of CNA but is unable to trigger its developmental program (**Fig. 4**).

We hypothesized that a small number of amino acid changes may be sufficient to convert the CNA-KnC3HDZ chimera into one with regulatory properties more closely resembling CNA. To identify these amino acids, we took advantage of the fact that CNA ortholog START domains displayed differential substitutability in our molecular and phenotypic assays (**Figs. 4a,b**). Using multiple sequence alignments, we found eight residues that are conserved only in fully-substituting chimeras (**Extended Data** Fig. 12). These residues were then introduced into the START domain of CNA-KnC3HDZ to create CNA-KnC3HDZ-8m. Supporting the functional relevance of these choices, a *pREV:CNA*-KnC3HDZ-8m* transgene conditioned partial HD-ZIPIII gain-of-function phenotypes in primary transformants whereas the *pREV:CNA*-KnC3HDZ* transgene did not (**Extended Data** Fig. 13).

To test the effects of the eight substitutions on target selection, we identified genomic regions bound by CNA-KnC3HDZ and CNA-KnC3HDZ-8m using ChIP-seq. CNA-KnC3HDZ bound 8,480 sites in the genome corresponding to 11,850 bound genes, and its binding profile is fully contained within that of CNA (**Figs. 6a-c**). By contrast, CNA-KnC3HDZ-8m showed a slightly more expanded binding profile, recognizing 8,871 sites in the genome, corresponding to 12,228 bound genes (**Figs. 6a-c**). Interestingly, 26% of all sites newly recognized by CNA-KnC3HDZ-8m (102 out of 392) are shared with CNA (**Figs. 6a-c**). Thus, eight amino acid substitutions can shift the binding profile of CNA-KnC3HDZ towards that of CNA.

**Figure 6.**
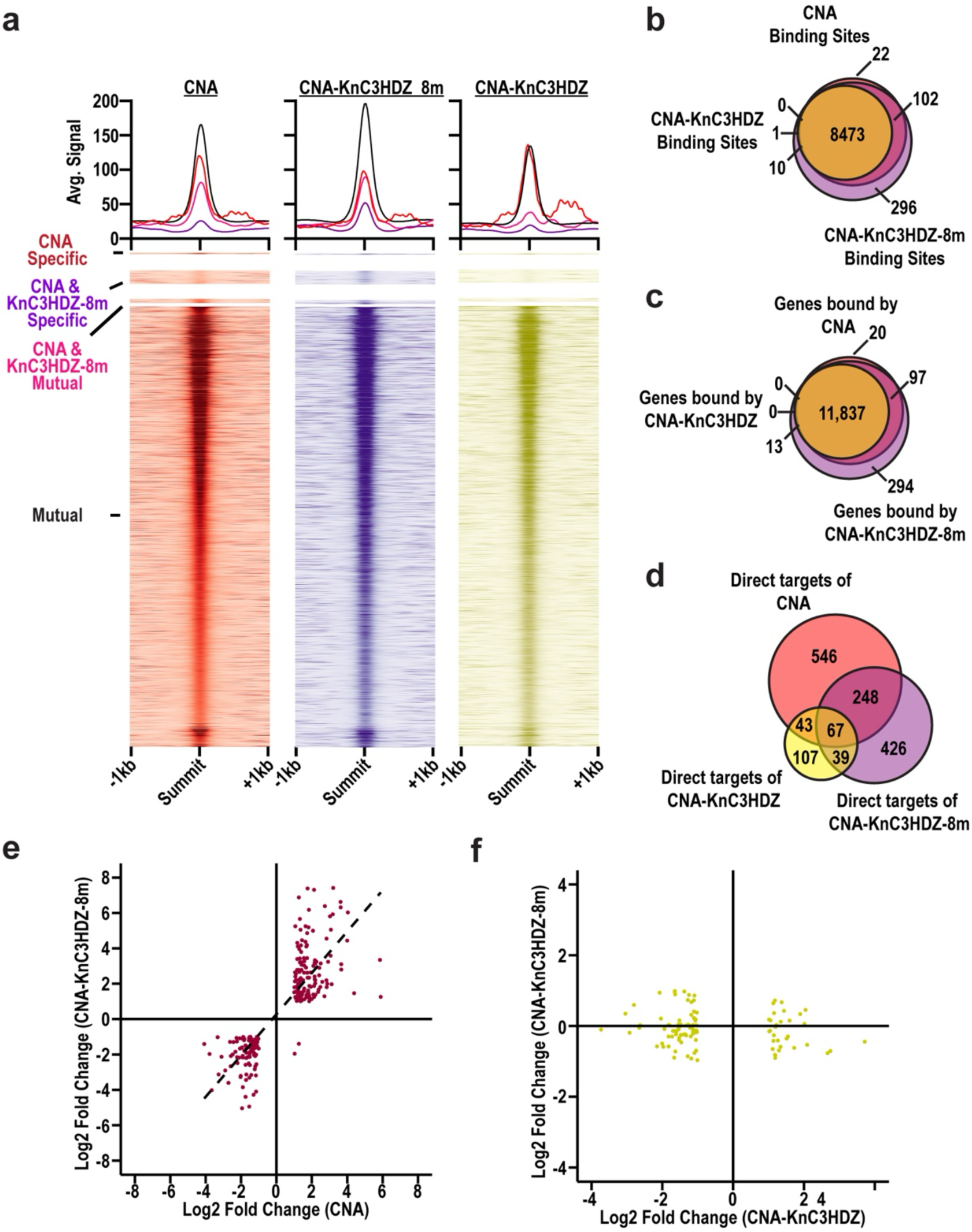
Evolutionarily relevant START point mutations alter target selection and regulation. **a,** Histograms and heatmaps of ChIP-seq signal intensities. Histogram lines and colors delineate four categories of binding sites: CNA specifically bound (red, solid); CNA-KnC3HDZ_8m specifically bound (purple, solid); bound by CNA and CNA-KnC3HDZ_8m (pink, solid); and bound by all three variants (black, solid). Heatmaps are separated into the same four categories. **b**, Venn diagram of sites bound by CNA and its chimeras. **c,** Venn diagram of genes bound by CNA and its chimeras. **d,** Venn diagram of direct targets of CNA and its chimeras, i.e. bound in ChIP-seq and differentially expressed in RNA-seq. **e,** Scatterplot showing genes bound by all three variants whose expression is regulated only by CNA and CNA-KnC3HDZ_8m (purple). **f**, Scatterplot showing genes bound by all three variants whose expression is regulated only by CNA-KnC3HDZ (yellow).

To test the effects of the eight substitutions on target regulation, we identified direct targets of CNA-KnC3HDZ and CNA-KnC3HDZ-8m by pairing ChIP-seq and transcriptome profiling. 378 and 1,151 genes were differentially expressed in KnC3HDZ- and KnC3HDZ-8m-expressing lines, corresponding to 256 and 780 high-confidence direct targets, respectively (**Fig. 6d; Supp Table 3g,h**). Of the 248 direct targets shared only by CNA and KnC3HDZ-8m, 7 were from genes newly recognized by KnC3HDZ-8m. This supports the notion that selection and regulation of new loci is not the primary mechanism by which START domains generate TF-specific outcomes. The remaining 241 direct targets are derived from loci that are bound by all three proteins, but only regulated by CNA and KnC3HDZ-8m (**Fig. 6e; Extended Data** Fig. 8). These direct targets correspond to genes with sites considered non-responsive by CNA-KnC3HDZ which are now considered responsive by CNA-KnC3HDZ-8m. Finally, KnC3HDZ has 107 unique direct targets, despite its binding profile being fully contained within that of CNA and KnC3HDZ-8m (**Fig. 6f; Extended Data** Fig. 8). These direct targets correspond to genes with sites considered responsive by CNA-KnC3HDZ which are now considered non-responsive by CNA-KnC3HDZ-8m. The latter two findings support the notion that START domains mediate the interconversion of responsive and non-responsive binding sites. Taken together, these developmental and functional genomics assays demonstrate that a small number of amino acid substitutions can lead to large-scale changes in the selection and regulation of HD-ZIPIII targets.

### Functional divergence of HD-ZIPIII proteins is partly explained by their START domains

Our findings indicate amino acid substitutions in HD-ZIPIII START domains lead to profound changes in target regulation. This supports the idea that START domains may contribute, at least in part, to HD-ZIPIII functional divergence. To formally test this notion, we repurposed the established complementation assay described in ^25^. In this assay, coding sequences of HD-ZIPIII genes were placed downstream of *REV* regulatory elements, introduced into the *rev-6* mutant^65^, and scored for their ability to complement the *rev-6* mutant phenotype^25^. Approximately 40% of *rev-6* mutant flowers terminate before setting seed^65^ (**Figs. 7a,b**), and this phenotype can be fully rescued by the introduction of a *pREV:REV* transgene^65^ (**Figs. 7a,b**). By contrast, *pREV:CNA* primary transformants are indistinguishable from *rev-6* mutants consistent with functional divergence of CNA from REV^25^ (**Figs. 7a,b**). We then replaced the CNA START domain with its counterpart from REV to create a CNA-AtREV chimera driven by *REV* regulatory elements (*pREV:CNA-AtREV*). Introduction of the *pREV:CNA-AtREV* transgene dramatically reduced the frequency of floral termination in *rev-6* mutants, with only 8% of siliques aborting during development (**Figs. 7a,b**). Thus, functional divergence of CNA and REV subclade members is driven, at least in part, by substitutions within their START domains.

**Figure 7.**
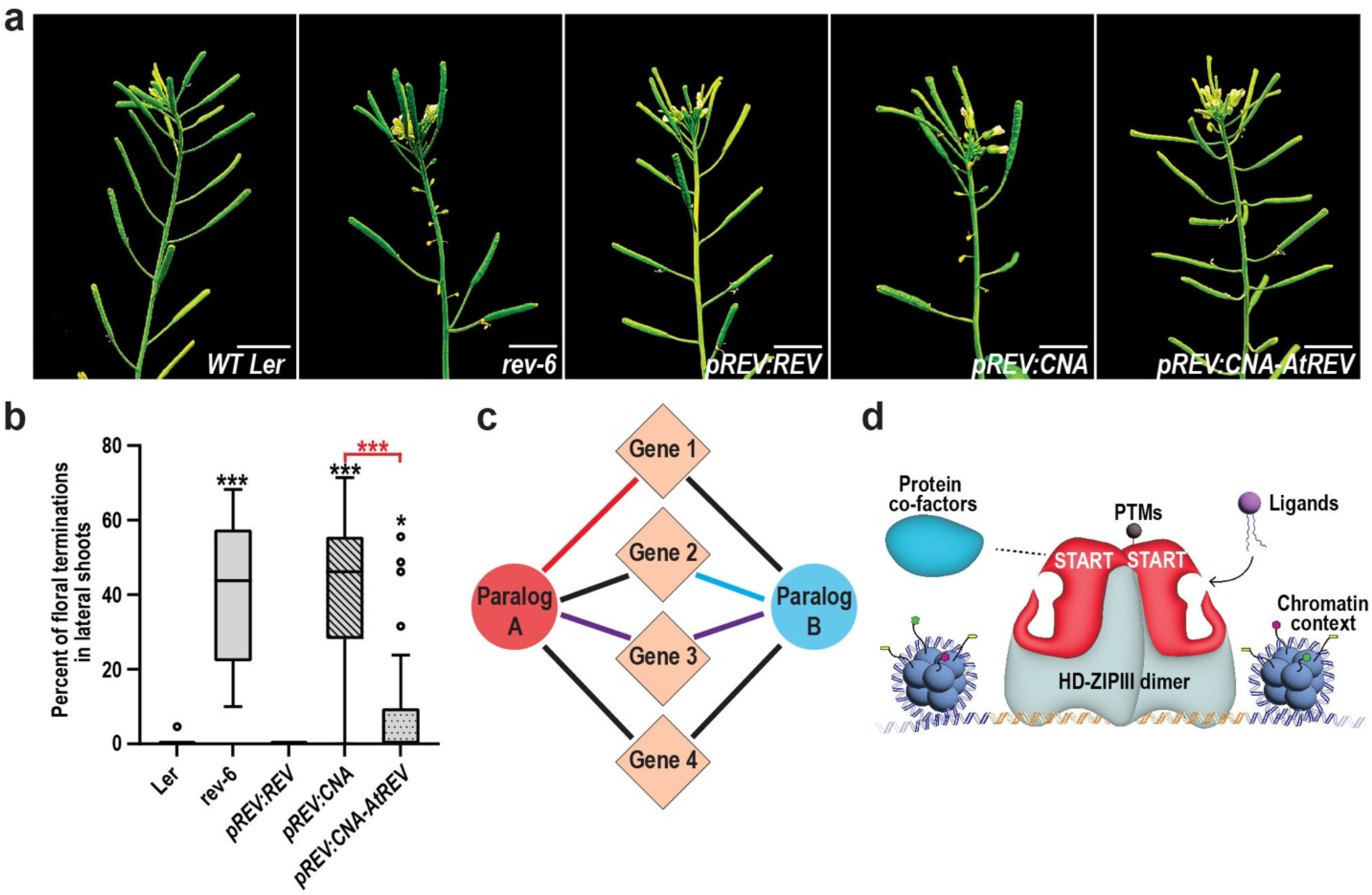
Functional divergence of HD-ZIPIII proteins is driven by their START domains which convert a single network architecture into paralog-specific regulons. **a**, The *rev-6* mutant has a strong floral termination phenotype^65^. The *pREV:REV* transgene fully complements the *rev-6* mutant phenotype as plants appear wildtype (L*er*), whereas the *pREV:CNA* transgene does not. The majority of primary transformants carrying a *pREV:CNA-AtREV* transgene appear wildtype indicating near-complete rescue of the *rev-6* mutant phenotype. **b**, Quantification of *rev-6* complementation (measured as percentage of floral termination in lateral shoots). Statistics are against the L*er* wildtype in all cases except the red asterisks which are compared to *pREV:CNA*. *** = p-value ≤ 1e-10. **c**, HD-ZIPIII TF paralog divergence through differential regulation of mutually bound genes. In this simplified model, two members of a TF family bind a common set of genes. A given gene could be regulated by both TFs (i.e. Gene 3) or unaffected by both TFs (i.e. Gene 4). Alternatively, a gene could be uniquely regulated by one of the two TFs (Gene 1 vs Gene 2) generating paralog-specific regulatory outcomes from a commonly bound genetic network. Line legend: purple (regulated by both paralogs), black (regulated by neither paralog), red (regulated only by paralog A), blue (regulated only by paralog B). **d**, Speculative, non-mutually exclusive regulatory mechanisms occurring at shared binding sites (orange DNA). Inputs integrated via START domains could include ligands (purple), post-translational modifications (PTMs; brown), and protein co-factors such as other TFs, chromatin remodelers, or subunits of the general transcriptional machinery (blue). Functional consequences for gene expression may also depend on histone modifications, DNA methylation, degree of chromatin compaction, or other aspects of chromatin context.

## DISCUSSION

TF functional divergence drives biological processes ranging from speciation to cellular decision making^66,67^. A common mechanism underlying this divergence is binding to different sets of loci^1,2,3,4,5,6^. Here we show that functional divergence of the HD-ZIPIII paralogs CNA and PHB is instead driven primarily by differential usage of shared binding sites. CNA and PHB have largely overlapping binding profiles, yet each paralog has hundreds of uniquely regulated targets that affect distinct biological processes (**Figs. 1e,g**; **Extended Data** Fig. 7). CNA and PHB also have hundreds of shared, similarly regulated direct targets, consistent with the partial functional redundancy of these paralogs^25^ (**Fig. 1f**). Thus, regulation of a given gene depends on whether its local binding site is considered responsive versus non-responsive. Interestingly, the interpretation of binding site identity by HD-ZIPIII proteins appears to be controlled, at least in part, by their START domain. For instance, as few as eight amino acid substitutions in the START domain of a chimeric CNA protein were sufficient to drive both gain and loss of regulation across hundreds of bound loci (**Fig. 6**). In addition, replacing the CNA START domain with that of REV enabled CNA to partly fulfil the developmental functions of its divergent paralog (**Fig. 7b**). Our findings support a model in which HD-ZIPIII TFs use information integrated via their START domain to generate paralog-specific transcriptional outcomes across a network of commonly bound genes (**Fig. 7c**).

Binding of shared sets of genes by functionally divergent paralogous TFs has been noted previously. For instance, E2F TFs oppositely regulate a common set of cell-cycle related genes^112^, while MyoD and Myf5 induce histone modifications and recruit transcriptional machinery, respectively, at shared skeletal muscle specification and differentiation genes^113^. However, unlike the HD-ZIPIII family, the activities of these metazoan TFs are spatiotemporally separated by sequential expression dynamics^112,113^. The plant paralogs FT and TFL1, on the other hand, do spatiotemporally overlap and share a network of commonly bound genes, much like HD-ZIPIII TFs^22^. However, these TFs are thought to function by changing the valence of gene expression rather than making paralog-specific decisions on whether a given gene is to be regulated or not. The HD-ZIPIII family is a particularly clear example of the latter given their dramatic overlap in genomic occupancy. However, this mechanism is likely at play in other TF families as well, including those whose functional divergence was previously attributed solely to binding of new targets. Reanalysis of shared binding sites from the perspective of ‘responsive versus non-responsive’ is likely to yield additional, biologically relevant insights. Related to this, genomes are characterized by prevalent ‘non-functional DNA binding events’, prompting animal transcription networks to be described as “Continuous Networks” (reviewed in ^53^). In this model, TFs occupy broad swathes of the genome, and occupancy levels positively correlate with functional consequences for gene expression. However, our analyses indicate occupancy levels are largely uncoupled from target regulation and magnitude of gene expression changes (**Fig. 1; Extended Data** Figs. 4a, 4b, 5a, 6). Thus, broad occupancy followed by functional discrimination of responsive versus non-responsive sites, may be a more complete description of how TFs engage with genomes.

One advantage of paralogs binding to non-overlapping, discrete sets of loci is that it minimizes inappropriate cross-regulation. What advantages might come from an alternate strategy in which a family of TFs occupies the same genes but regulates transcriptional outcomes on a paralog-specific basis? One possibility is suggested by the opposite phenotypes of the *phb phv rev* and *phb phv cna* triple mutants^25^. The former has a meristem-termination defect while the latter has enlarged fasciated meristems^25,68^. These phenotypes only emerge when both *PHB* and *PHV* are mutated, suggesting PHB and PHV buffer the meristem-promotive effects of REV and the meristem-repressive effects of CNA. One mechanism by which PHB and PHV could accomplish this buffering is occupation – but not necessarily regulation – of the genes REV and CNA use to affect meristem size. In this scenario, PHB and PHV would prevent meristem termination in a *rev* single mutant by limiting the effects of the meristem-repressive CNA. Similarly, in a *cna* single mutant, PHB and PHV would prevent meristem enlargement by blocking runaway effects of the meristem-promotive REV. This strategy would lend robustness to the transcriptional outcomes of each paralog and therefore the developmental contributions of the family.

Another possible advantage could be ease of network rewiring. For instance, it may be useful for a TF paralog to gain or lose control of entire regulatory modules, or for a diverged paralog to regain certain targets given environmental or evolutionary pressures. This would be mechanistically challenging for paralogs which diverged *via* changes to genome occupancy. By contrast, a strategy of occupation – but not necessarily regulation – of target genes preserves connections between members of a TF family and their downstream targets (**Fig. 7c**). Inputs from respective paralogs can then be turned on or off in a coordinated fashion. This could be accomplished by stimuli operating at organismal or population levels, increasing speed and flexibility of rewiring. Paralog-specific stimuli could then be used to further expand regulatory potential within a given TF family.

The acquisition of additional protein domains by homeobox TFs is thought to have provided new targets and increased specificity of binding^69^. Consistent with this, our findings indicate a weak relationship between START domains and the frequency of HD-ZIPIII genomic occupancy, perhaps accomplished by tuning of DNA binding affinities (**Figs. 1,3,5,6**). More importantly, our analyses suggest that capture of START domains by HD-ZIPIII precursors added an additional layer of regulation at the level of binding site usage. How might differential usage of shared binding sites occur? CNA and PHB have nearly identical endogenous expression patterns^70^, and the estradiol-inducible system generates uniform TF accumulation in all cell types^48^ (**Extended Data** Fig. 1). This argues against spatiotemporal separation of requisite co-factor(s) as a mechanistic explanation. However, formally ruling this out will require single cell resolution assays that control for heterogeneity inherent to whole tissue experiments. Instead, one potential explanation for distinct START-directed licensing of CNA and PHB could be paralog-specific ligands. For instance, the PHB START domain binds to phosphatidylcholine, and this binding is required for PHB to occupy *ZPR* regulatory regions and strongly activate their transcription^48^. Binding of other ligands could conceivably uncouple these two features, creating situations in which a paralog binds to a given gene but cannot affect its transcription. A second non-mutually exclusive mechanism to facilitate the interpretation of binding site identity is paralog-specific interacting partners. These partners could directly interact with the START domain, as observed for other StARkin-containing proteins^41,44,62,71,72,73,74,75^. Co-factor binding could also occur at other regions of the HD-ZIPIII protein and be modulated by START-directed allosteric effects. In either scenario, the accrual of amino acid substitutions throughout evolution would lead to distinct interactomes for each paralog. Interactions could then be used to convert non-responsive binding sites into responsive ones and vice versa. Numerous mechanisms could mediate this interconversion including opening or closing of chromatin, formation of higher-order chromatin structures, and recruitment and/or stabilization of basal transcriptional machinery, among others.

We note that StARkin domains use a remarkably diverse array of regulatory mechanisms and are present in gene families across the tree of life^45,58,59,60,61,62,77,78^. Their ubiquity and flexible regulatory nature may make StARkin domains particularly amenable tools for evolution to generate functional divergence. Studies into other StARkin-containing multigene families will be instrumental in testing this intriguing hypothesis.

Finally, we mention some caveats of the work presented here. First, ectopic inducible systems have numerous advantages. For instance, they can circumvent confounding morphological defects of mutants or stable overexpression lines, standardize TF dosage, and eliminate spatiotemporal variance of co-factors. However, supplementary experiments using endogenous promoters are critical for findings to be properly integrated back into a developmental context. Second, pairing ChIP-seq with transcriptome profiling is an efficient approach to identify direct targets (e.g. ^111^). However, these assays can suffer from false positives (Type I errors) and false negatives (Type II errors), influenced in part by induction time prior to transcriptome profiling. Shorter inductions capture strongly activated targets but are more likely to miss repressed targets that require additional time to be detected. By contrast, longer inductions are better positioned to capture both activated and repressed targets but will miss genes whose regulation has been dampened by network feedback. Combining both time frames is a good strategy to capture the full range of likely TF direct targets.

## METHODS

### Arabidopsis Growth Conditions

Arabidopsis thaliana (Col-0 and Ler-0 ecotype) seedlings were grown at 22°C under long-day conditions on soil or 1% agarose Murashige and Skoog medium plates (pH 5.7). Inductions were performed by spraying 9 day old T2 seedlings on agarose plates 10 times with a 50uM β-Estradiol, 1% DMSO, and 0.005% Silwet L-77 solution.

### Molecular cloning and plant transformation

PHB constructs (*pOlexA:PHB* & pOlexA:PHB*-SDΔ*) used in this study had been previously generated^48^. *CNA-mCitrine-3xFLAG* was constructed via Gibson assembly (NEB) in a pCR8/GW/TOPO cloning plasmid backbone. The *CNA-SDΔ* CDS was constructed via Gibson assembly with primers that remove the START domain (451 – 1149 bp from start codon) but retain the 21-nt regulatory miR166 binding site, GGAATGAAGCCTGGACCGGAT (553 – 573 bp from start codon) through overlapping primers. To generate the *CNA-SDmut* CDS, the mutated variant of the START domain was synthesized (GeneArt), replacing the amino acid sequences RDFWLLR and RAEML to GAVVGVG and VAAGV, respectively^48^ using the following nucleotide substitutions: CGCGATTTCTGGCTGTTACGT (802 – 822 bp from start codon) to GGTGCCGTCGTAGGAGCAGGC and AGAGCAGAGATGCTT (922 – 936 bp from start codon) to GTGGCGGCCGGCGTC. miR166 insensitivity (*CNA*, CNA-SDmut*, and CNA-SDΔ\**) was introduced by PCR amplification of the START domain in two fragments using mismatched primers to introduce a silent SNP within the 21nt miR166 recognition site (GGAATGAAGCCTGG**<u>T</u>**CCGGAT to GGAATGAAGCCTGG**<u>A</u>**CCGGAT, 553 – 573 bp from start codon) followed by reassembly via Gibson assembly.

CNA START domain substitutions were generated via Gibson assembly by amplifying the CNA CDS and pCR8 backbone to exclude the START domain (451 – 1149 bp from start codon) and insertion of orthologous CNA START domains made miR166 insensitive, if necessary, as previously described from cDNA (KnC3HDZ, PpHOX32, SmHOX32, ZmHOX29, AtREV) or via gene synthesis (CaC3HDZ, CharoAnc, AngioAnc). Inducible pOlexA constructs were generated by subcloning into the two component system (pUBQ:XVE pOlexA:CDS; modified pMDC7) binary vector^48^ via LR Gateway reaction. Constructs with the regulatory elements of CNA (pCNA, 3728 bp upsteam^63^ ) or REV (pREV, 4975 bp upsteam of the start codon upsteam, 1026 bp downstream of stop codon) were assembled via Gibson assembly and subcloned into a pB7GW binary vector. *A. thaliana* was transformed with these constructs via *Agrobacterium* mediated floral dip transformation. Cloning primers and synthesized sequences can be found in **Supp. Table 4a,b**.

### Confocal imaging

Induced seedlings were incubated in a 0.5µg/ml propidium iodide solution for 5 minutes to stain root cell plasma membranes. Seedlings were washed 3 times for 5 minutes each with dH_2_O. The hypocotyl was excised, and roots were wet mounted to a glass slide with coverslip. Roots were examined under a Nikon Eclipse Ni-E microscope with C2si confocal system using a 60x Nikon Plan APO VC 60x / 1.40 oil immersion objective lens with Cargille Type A immersion oil. The mCitrine YFP tag was excited using a wavelength of 526nm, and emission was captured between 490 – 550nm. Propidium iodide was excited with the same wavelength as mCitrine, 526nm, and emission was captured between 570 – 620nm.

### Chromatin Immunoprecipitation and qPCR

Induced seedlings (24hrs post-induction) were collected off the plates and moved to a conical tube with 30ml of a 2% formaldehyde (ThermoFisher catalog# 28906) crosslinking solution. Uncapped tubes were placed in a vacuum chamber for 30 minutes at 20-25 mmHg vacuum pressure, with a brief shake after the first 15 minutes. The crosslinking reaction was inactivated by adding 2ml of 2M glycine to the formaldehyde solution, briefly shaking, and placing the uncapped tubes into the vacuum for 5 additional minutes. The inactivated formaldehyde solution was poured out and the seedlings were rinsed three times with 1xPBS. Paper towels were used to dry the seedling, removing as much excess water as possible. The dried seedlings were separated into individual 500mg biological replicates and flash frozen with liquid nitrogen.

Seedlings were ground using a mortar and pestle on dry ice, and the powdered tissue was transferred to a conical tube containing 8ml of nuclear isolation buffer (10mM HEPES pH 8.0, 1M Sucrose, 5mM KCl, 5mM MgCl_2_, 0.6% Triton, 0.4mM PMSF, 1x cOmplete™ EDTA free protease inhibitor) and rotated for 15 minutes at 4°C. The solution was filtered through two layers of Miracloth into a separate conical tube and centrifuged at 3000g for 15 minutes at 4°C. The supernatant was discarded, and the pelleted nuclei were resuspended in 1ml of nuclei wash buffer (10mM Tris pH 8.0, 250mM Sucrose, 10mM MgCl_2_, 1mM EDTA, 1% Triton, 1mM PMSF, 1x cOmplete™ EDTA free protease inhibitor), transferred to a microcentrifuge tube, and centrifuged at 12,000g for 10 minutes at 4°C. The supernatant was discarded, and the nuclei were resuspended in nuclear lysis buffer (20mM Tris pH 8.0, 2mM EDTA, 0.01% SDS, 1mM PMSF, 1x cOmplete™ EDTA free protease inhibitor) and transferred to a 1ml Covaris milliTUBE with AFA Fiber. Samples were sonicated using a Covaris E220 at 150W peak power, 20% duty factor, 200 cycles/burst for 6 minutes. The sonicated chromatin was transferred to a new microcentrifuge tube and centrifuged at 12,000g for 10 minutes at 4°C to pellet insoluble debris. 920ul of the supernatant was moved to a new microcentrifuge tube, and 30ul of 5M NaCl and 20ul of 30% Triton were added to the supernatant. The sonicated chromatin was divided into two microcentrifuge tubes (420ul each), and 42ul of the remaining chromatin was retained for a 10% input control. 2ul of M2 mouse monoclonal anti-FLAG (Sigma, F3165) or mouse IgG was added to the tubes and rotated overnight at 4°C. Pierce Protein A/G magnetic beads (Thermofisher Scientific, 88802), were equilibrated with the nuclear lysis buffer with added NaCl and Triton as described previously and resuspended to their initial volume 40ul of equilibrated magnetic beads were added to each tube, and incubated for 2hrs rotating at 4°C. Tubes were then placed on a magnetic rack and the supernatant was removed. Beads were washed twice for 5 minutes each rotating at 4°C with the subsequent buffers, Low Salt Wash Buffer (150mM NaCl, 0.1% SDS, 1% Triton x-100, 2mM EDTA, 20mM Tris pH 8.0), High Salt Wash Buffer (500mM NaCl, 0.1% SDS, 1% Triton x-100, 2mM EDTA, 20mM Tris pH 8.0), LiCl Wash Buffer (250mM LiCl, 1% IGEPAL, 0,1% SDS, 1mM EDTA, 10mM Tris pH 8.0) and TE Buffer (10mM Tris pH 8.0, 1mM EDTA, 0,1% IGEPAL). 250ul of preheated (65°C) Elution Buffer (1% SDS, 0.1M NaHCO_3_) was added to the beads and 10% input and incubated at 65°C for 15 minutes. The supernatant was transferred to a new microcentrifuge tube, 10ul of 5M NaCl was added to each tube, and placed at 65°C overnight. The heat block was reduced to 45°C and 10ul of 0.5M EDTA, 20ul 1M Tris pH 7.0, 1ul of 20mg/ml Protease K, and 1ul Rnase (50ng/ul) were added to each to and incubated for 1 hour at 45°C. ChIP DNA was recovered using a Zymo DNA Clean and Concentrator-25 kit.

Recovered DNA was quantified using qPCR with 10ul of SybrGreen, 1ul of combined 10uM forward and reverse primer sets, 5ul dH2O, and 4ul of DNA with two technical replicates for each tested target (*OTC*, *ZPR3*, and *ZPR4*). Data was plotted using GraphPad.

### ChIP-seq and bioinformatics

Libraries were constructed using the NEBNext Ultra II DNA Library kit and single-end sequenced using the Illumina NextSeq500/550 located on-site at the Penn Epigenetics Institute. Read files were merged using command prompt in Windows OS, using the command: “Type file_L001.fastq.gz file_L002.fastq.gz file_L003.fastq.gz file_L004.fastq.gz > merged-file.fastq.gz”; if necessary and uploaded to the Galaxy web-based platform^80^.Reads were trimmed using Trimmomatic^81^ and performing the initial ILLUMINACLIP using adaptor sequences for TruSeq3 (single-ended, for MiSeq and HiSeq) and the trimmomatic operation “Sliding window trimming” averaging across 4 bases with an average required score of 20. Trimmed sequencing files were aligned to the TAIR10 Arabidopsis thaliana genome using Bowtie2^82^ under the default settings. Number of mapped read statistics can be found in **Supp. Table 5a.**

Peaks were called using MACS2^83^ running the individual FLAG IP samples individually against their respective 10% input control, using the effective genome size of *D. melanogaster* equal to 1.2e8 (equivalent to the genome size of *A. thaliana*) and building a shifting model with a lower mfold bound = 5, upper mfold bound = 50, and a band width for picking regions to compute fragment size = 300, and the minimum FDR (q-value) cutoff for peak detection set to 0.01. Window size was set to 150bp then BED files were analyzed using a publicly available Irreproducibility Discovery Rate (IDR) python package (https://github.com/nboley/idr). An IDR of 0.01 was used (following ChIP-Hub guidelines for plant genomic analyses^114^), and we kept only the peaks that were called in all replicates of a given genotype. Diffbind^84^ was then used to analyze differential peak signals between samples, and further analyzed with custom R scripts to perform a 3-way comparisons of DiffBind results and additional outputs to include bound genes using the TAIR10 gff3 genome annotation with further integration of DEseq2 RNA-seq results (Supp. Tables 2,3) using R packages profileplyr^85^, dplyr^86^ , reshape^87^, stringr^88^, and rtracklayer^89^. A peak was assigned to a gene if it was <2kb upstream of its transcription start site, within the gene body, or <1kb downstream of its transcription termination site. Histograms were plotted using GraphPad (Prism) and heatmaps were generated using ggplot2^90^.

### MEME-ChIP motif identification

BED coordinates of the binding sites centralized on the summits for each transcription factor were uploaded as custom tracks to the UCSC genome browser^91^ and the TAIR10 genome. Using Table Browser feature^92^, DNA sequences in fasta format of these 400bp summits, and an additional 50bp upstream and downstream (total 500bp centralized on the summit) were extracted. There sequences were then uploaded to the MEME-ChIP discovery tool^93^ of the MEME suite 5.5.5^94^. The “Input the motifs” setting was set to known motifs of “ARABIDIPSIS (Arabidopsis thaliana) DNA” and “DAP motifs^49^”. The 2^nd^ order model of sequences was used as the background model, looking for motifs with widths between 6 and 11 in MEME and STREME. Zero or one occurrence per sequence was selected for the expected motif site distribution and MEME was restricted to search for palindromes only.

### FIMO identification of bound motifs and DNA shape prediction

Individual instances of the top motif for CNA and PHB mutual sites (VTAATNATTAB for CNA; TAATRATKATD for PHB) were identified using FIMO^95^ in the MEME suite 5.5.5^94^, with a p-value cutoff of 5e-4 and input sequences of Ensemble Plants Genomes and Proteins: Arabidopsis thaliana (version 57). The output bed file of genomic coordinates with this motif were uploaded as a custom track to UCSC genome browser^91^ and the table browser tool^92^ was used to find any overlap of these motif genome coordinates with regions 75bp upstream or downstream from the summits of mutual CNA and PHB peaks.

Table browser was used to output sequences of the bound and unbound motifs in the genome with an addition 8bp upstream and downstream. Fasta sequences were then converted into DNA shapes using the DNAshapeR package^96^ in R. Two-tailed Z-tests were then performed for predicted minor groove width (MGW), helical twist (HelT), propeller twist (ProT) and roll at individual position along the sequences between the shapes of bound and unbound motifs and flanking sequences, and q-value was calculated using Bonferroni correction^97^).

### RNA-seq and bioinformatics

All seedlings were induced (or mock induced) two hours after the beginning of the light cycle to minimize circadian differences between genotypes. Seedlings were then collected 24hrs post treatment and flash frozen in liquid nitrogen. Total RNA was extracted using TRIzol Reagent (Thermofisher Scientific, 15596026) and treated with DNAse. Sequencing was performed by Lexogen (Austria) using the QuantSeq 3’ mRNA protocol which prioritizes the 3’ end of the mRNA transcripts. Sequencing files were demultiplexed by Lexogen, and reads were uploaded to the Galaxy web-based platform^80^. Adaptor sequences were trimmed using Trimmomatic^81^, performing the initial ILLUMINACLIP using adaptor sequences for TruSeq3 (single-ended, for MiSeq and HiSeq) and the trimmomatic operation “Sliding window trimming” averaging across 4 bases with an average required score of 20. Remaining polyA tail sequences were removed with a subsequent rerun of trimmomatic, performing an initial ILLUMINACLIP with custom adaptor (PolyA20) sequences of “AAAAAAAAAAAAAAAAAAAA” and the trimmomatic operation “Sliding window trimming” averaging across 4 bases with an average required score of 20.

Reads were aligned to the TAIR10 *A. thaliana* reference genome using RNA STAR^98^ with the following settings: Length of the SA pre-indexing string = 12; Maximum number of alignments to output a read’s alignment results, plus 1 = 200; Minimum overhang for spliced alignments = 8; Minimum overhang for annotated spliced alignments = 1; Maximum number of mismatches to output an alignment, plus 1 = 999; Maximum ratio of mismatches to mapped length = 0.6; Minimum intron size = 20; Maximum intron size = 1000; Maximum gap between two mates = 1000, Maximum number of collapsed junctions = 5000000. Number of mapped read statistics can be found in **Supp. Table 5b.**

Gene expression was then measured using featureCounts^99^ using the TAIR10 gene annotation gff3 and TAIR10 transcriptome using the following settings: Specify stand information = Forward; GFF Feature Type filter = gene; GFF gene identifier = ID; Exon-exon junctions = Count reads supporting each exon-exon junction (-J).

Results of featureCounts were combined into a single csv file and analyzed with DESeq2^100^ in R under the default settings. Comparison of respective induced and mock samples of a genotype were run through the lfcShrink function to account for the log fold change standard error across the samples using type = “ashr”^101^, shrinking the Log2 fold change values. Differentially expressed genes (DEGs) were determined using DESeq2, comparing induced genotypes to their respective mock control. Significant DEGs were identified as having ≥ 2 or ≤ -2fold change in expression levels, and a q-value ≤ 0.1. Significant DEGs were then referenced to bound genes identified as part of the ChIP-seq analysis to identify true targets. Data was plotted using the ggplot2 package^90^ in R.

### Gene Ontology Analysis

Gene ontology analysis was performed as in ^102^. In brief, a custom wrapper around the topGO package version 2.54.0^115^ was run on upregulated or downregulated direct targets for PHB specific, CNA specific and mutual targets. The gene universe was all detected transcripts in the RNA-Seq dataset. Packages were obtained from CRAN or Bioconductor version 3.7^116^. The complete output of gene ontology terms can be found in **Supp. Table 6**.

### qRT-PCR

cDNA was generated from 500ng total RNA from 24hr post-induced seedlings SuperScriptIV Reverse Transcriptase kit (ThermoFisher). cDNA was diluted 1:1 with dH2O and prepared in a 20ul volume (10ul of SsoAdvanced Universal SYBR Green Supermix (Bio-Rad), 8ul of dH2O, 1ul of cDNA, and 1ul of a 10uM mixture of the appropriate forward and reverse primer). Ct values were measured using a Eppendorf Mastercycler RealPlex^2^ with 40 cycles. Averaged Ct values of the technical replicates were normalized ACTIN 2 (ACT2) by subtracting the averaged Ct value of ACT2. Relative expression levels normalized to ACT2 were calculated with the formula: 2 ^(-1^ ^x^ ^normalized value)^. Data was plotted with Graphpad (Prism).

### Ancestral Sequence Reconstruction

Thirty CNA orthologs were identified using reciprocal best hit with BLASTp of the National Center for Biotechnology Information (NCBI) toolkits^103^ . These sequences consist of five eudicots, five monocots, four gymnosperms, four ferns, five bryophytes, three lycophytes, and four charophytes (**Extended Data** Fig. 10). Full length amino acid sequences were aligned using the ClustalW algorithm in MEGA X^104,105^ . Phylogeny was inferred by Bayesian Markov chain Monte Carlo (BMCMC) analysis using MrBayes (Version 3)^106,107^ and the maximum likelihood method with Jones-Taylor-Thornton (JTT, gamma distribution = 4) model^108^. Predictions were made with 100,000 generations, sampling every 100 and the first 250 samples discarded as burn-in, achieving a standard deviation of 0.0017. Bayesian posterior probability scores of 1.0 suggested high probability of inferring sequences at the nodes where ancestral sequences of interest were located. Ancestral sequences were predicted using the maximum likelihood method with JTT +4 gamma model in MEGAX. Coding sequences were assigned using the A. thaliana CNA coding sequences as a template, retaining the codons of conserved residues, and assigning codons for non-conserved residues. These sequences were ordered through GeneArt as synthesized DNA in vectors.

### Multiple Sequence Alignment to generate KfC3HDZ(8m)

CNA ortholog amino acid sequences were aligned using the ClustalW algorithm^104^ in the MEGA X software^105^ . Conserved aligned residues were identified using R and “msa” package^109^ from Bioconductor. A 70% consensus threshold score was used to identify conservation in 5 of the 8 aligned START domains, identifying residues conserved in the START domains of CNA, AngioAnc, ZmHOX29, PpHOX32, and CharoAnc, but not conserved in SmHOX32, KnC3HDZ, or CaC3HDZ.

### Accession Numbers

CNA orthologs for chimeras: *Zea mays* (ZmHOX29, XP_008655743.1), *Selaginella moellendorffii* (SmHOX32, XP_002961227.1), *Physcomitrella patens* (PpHOX32, XP_024399219.1)*, Klebsormidium nitens* (KnC3HDZ, GAQ87733.1), and *Chlorokybus atmophyticus* (CaC3HDZ, AZZW_21737^110^).

## DATA AVAILABILITY

All relevant data generated in this study are deposited to public repositories and are publicly released. Raw and processed data of ChIP-seq and RNA-seq data are deposited at the NCBI Gene Expression Omnibus (https://www.ncbi.nlm.nih.gov/geo/query/acc.cgi?acc=GSE253676). Relevant R codes are available at github.com/HusbandsLab.

## ACKNOWLEDGEMENTS

We are grateful to Dr. Ken Zaret, Dr. Doris Wagner, and Nicole Callery for thoughtful comments that have improved the manuscript. Work in the Husbands Lab is funded by grants from the National Science Foundation: #2039489 and #2310356 to A.Y.H.

## SUPPLEMENTARY INFORMATION

**Supplementary Tables 1-6**

**Supplementary Table 1**. **a**, FIMO output of instances of CNA and PHB top motif in the A. thaliana genome using a p-value cutoff of 1e-4. **b**, Genomic coordinates of PHB and CNA mutual binding sites summits with 75bp upstream and downstream. **c**, Summary table of bound and unbound motifs. **d**, Summary of DNA shape features and comparison of bound and unbound motifs. **Supplementary Table 2.** DiffBind results. **a**, Comparison of CNA, PHB, and Col-0. **b**, Comparison of CNA, CNA-SDΔ, and Col-0. **c**, Comparison of PHB, PHB-SDΔ, and Col-0. **d**, Comparison of CNA, CNA-SmHOX32, and Col-0. **e**, Comparison of CNA, CNA-PpHOX32, and Col-0. **f**, Comparison of CNA, CNA-KnC3HDZ, CNA-KnC3HDZ_8m, and Col-0. **Supplementary Table 3.** DESeq2 comparison of estradiol induced constructs and mock inductions. **a**, CNA. **b**, PHB. **d**, PHB-SDΔ. **e**, CNA-SmHOX32. **f**, CNA-PpHOX32. **g**, CNA-KnC3HDZ. **h**, CNA-KnC3HDZ_8m. **Supplementary Table 4.** Primers and synthesized DNA sequences. **a**, Cloning primers. **b**, synthesized START domain sequences. **c,** qPCR and qRT-PCR primers. **Supplementary Table 5.** Mapping statistics of ChIP-seq and RNA-seq reads. **a**, ChIP-seq. **b**, RNA-seq. **Supplementary Table 6.** Gene ontology analysis.

---

**Extended Data Figure 1.**
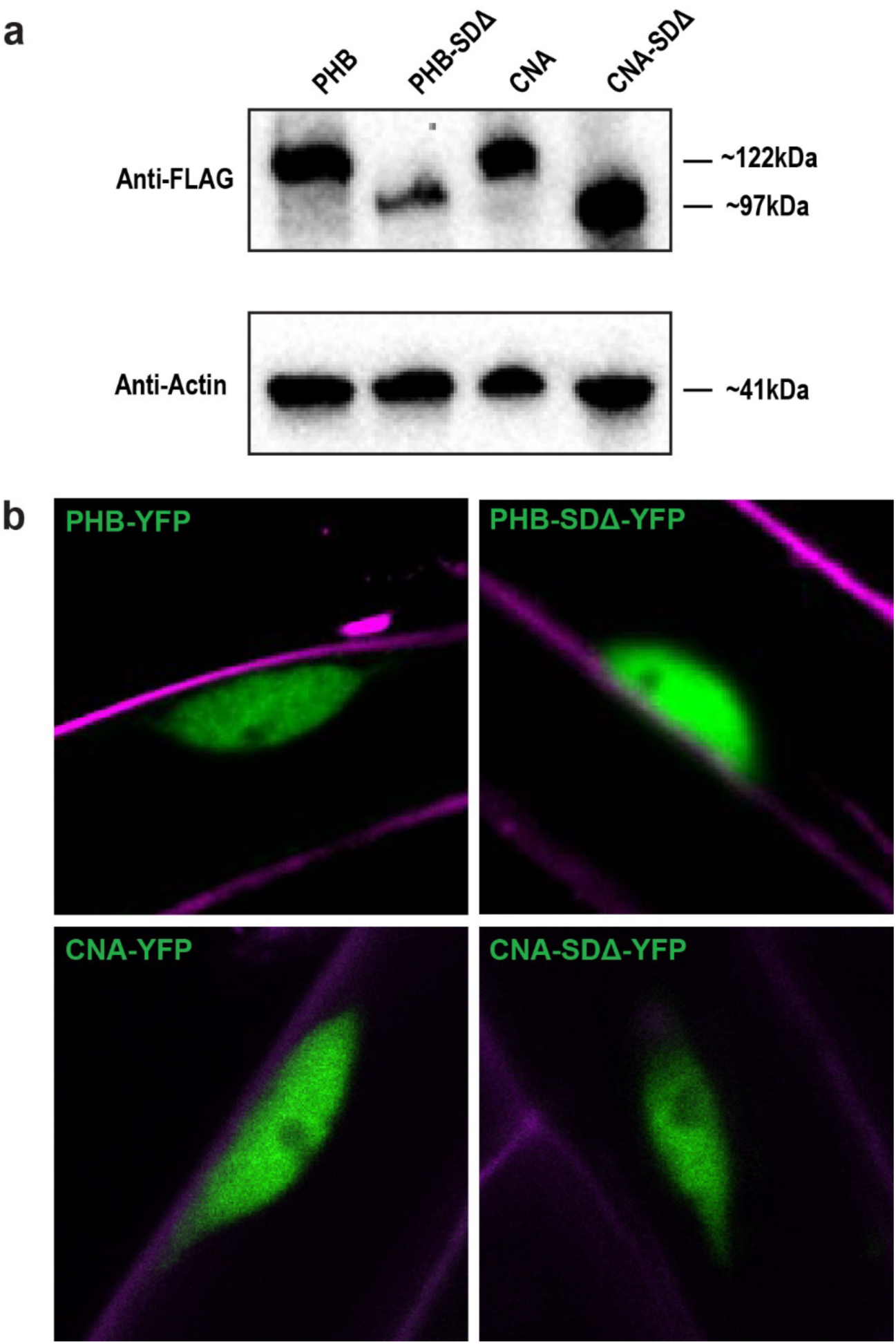
Estradiol-induction system controls. **a**, Western blots of induced pUBQ constructs showing robust induction of CNA and PHB variants in all transgenic lines. **b**, Confocal imaging of nuclei in estradiol-induced seedlings showing nuclear localization of all CNA and PHB variants (green). Tissue was stained with propidium iodide (magenta).

**Extended Data Figure 2.**
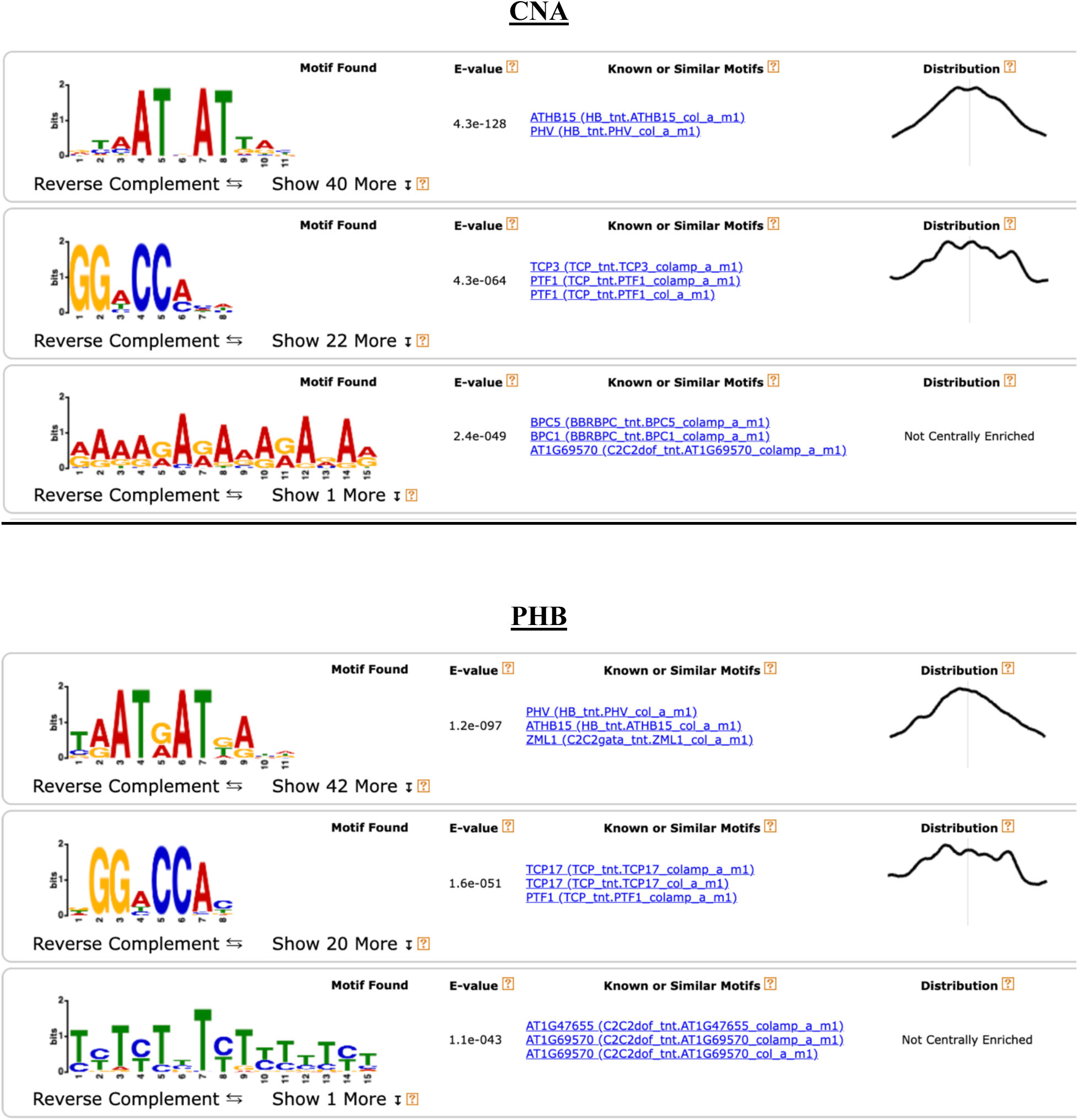
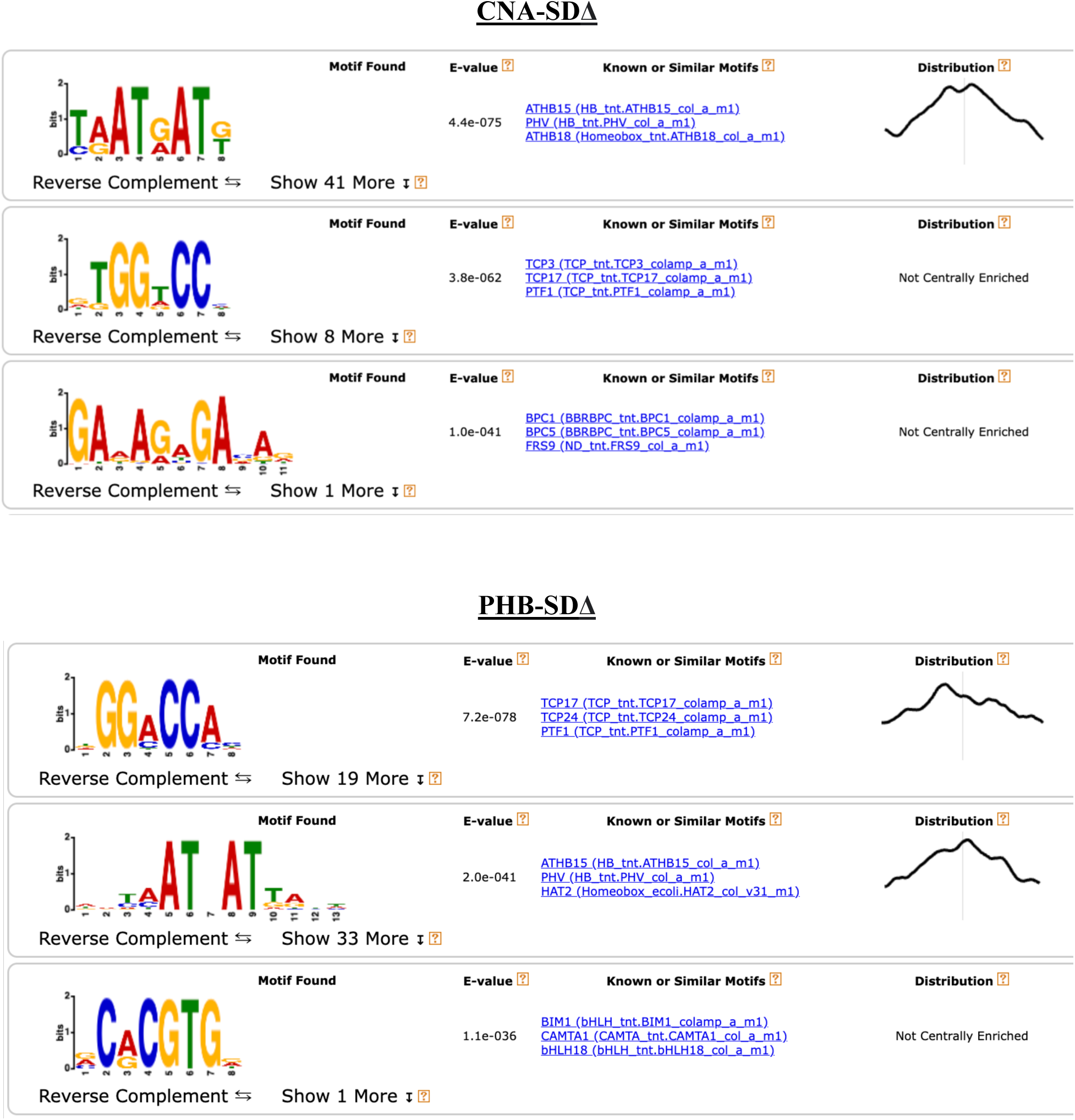
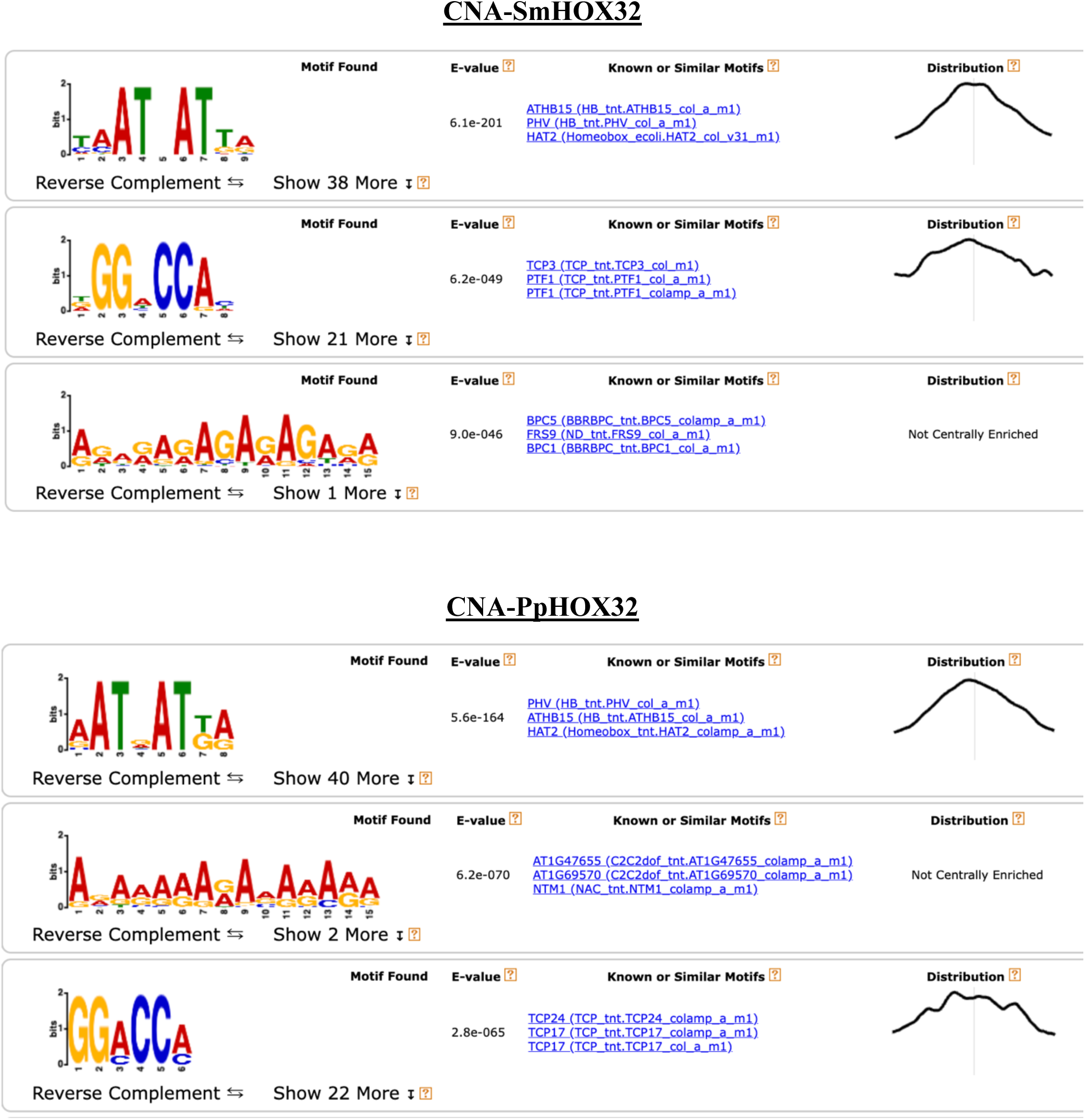
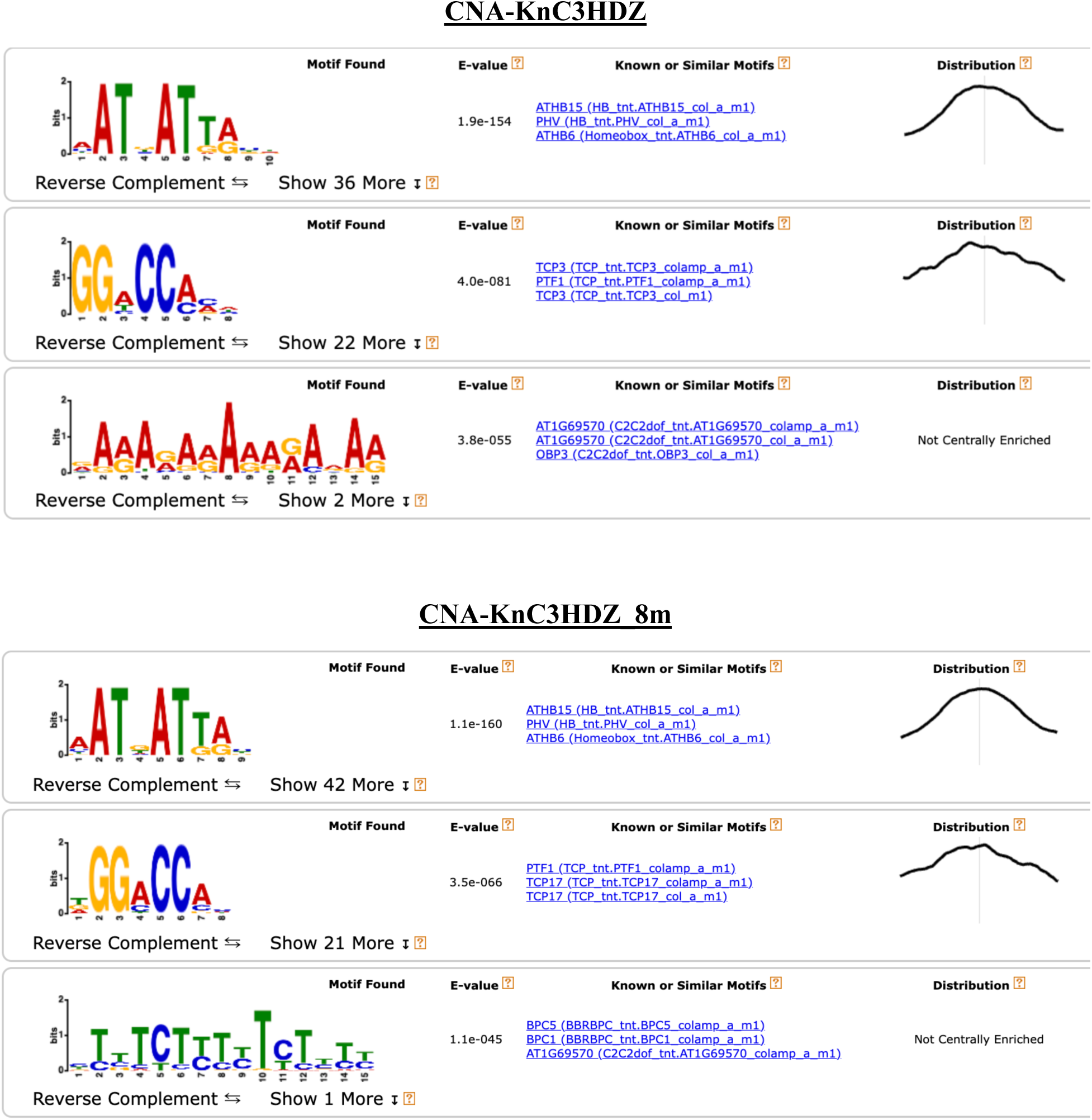
Top 3 motif results of MEME-ChIP of CNA and PHB binding sites. All variants bind an ATNATT consensus motif.

**Extended Data Figure 3.**
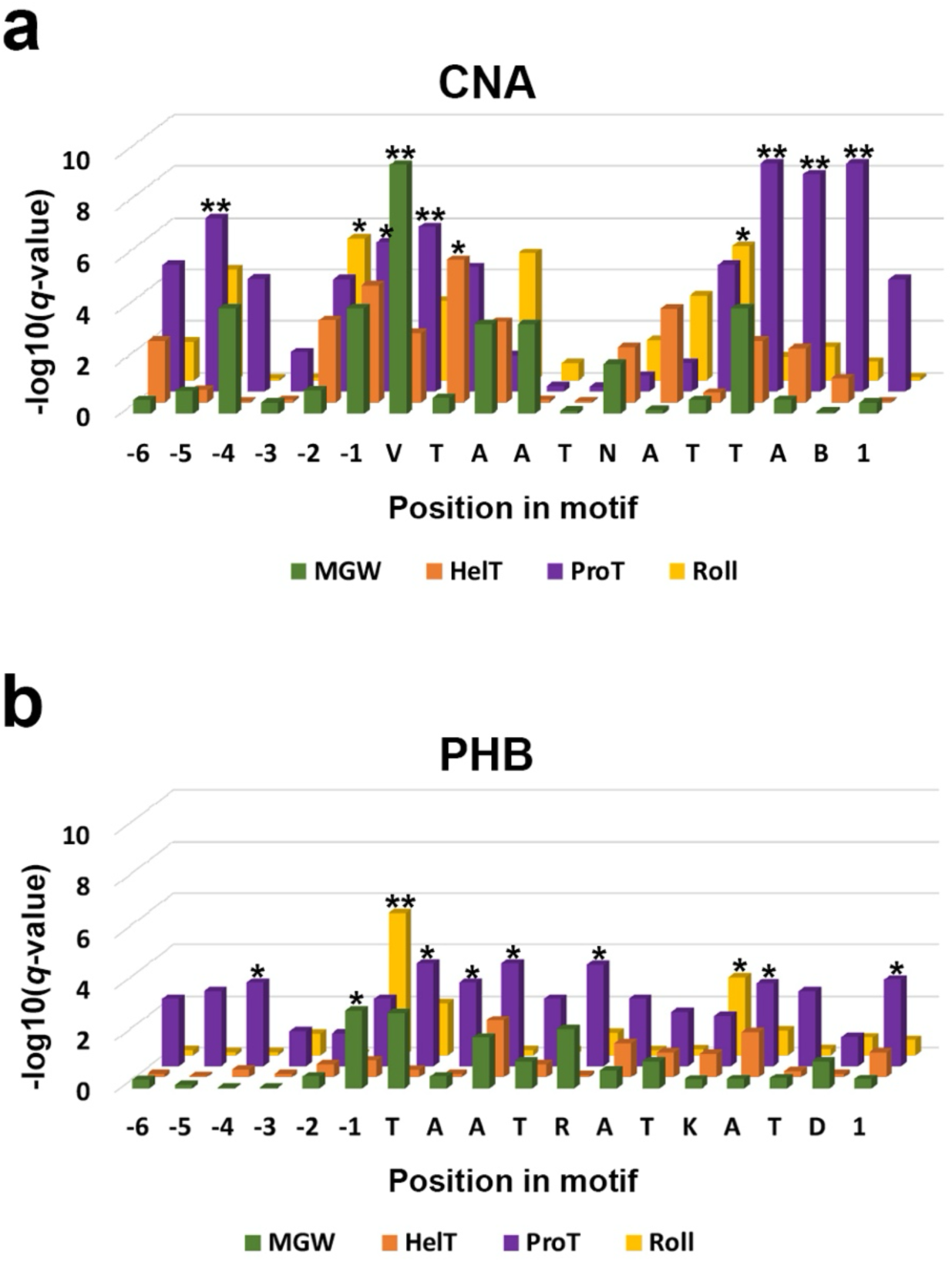
DNA shape features Z-test statistical differences between CNA bound, PHB bound, and unbound ATGATTA motif sequences in the *A. thaliana* genome. Motifs bound by CNA and PHB show strong deviations at multiple nucleotides across several axes of DNA shape. For clarity, only the most significant positions are marked with asterisks; see **Table S1** for *q*-values of all positions. CNA: * = *q*-value ≤ 10^-5^. ** = *q*-value ≤ 10^-6^; PHB: * = *q*-value ≤ 10^-3^. ** = *q*-value ≤ 10^-5^.

**Extended Data Figure 4.**
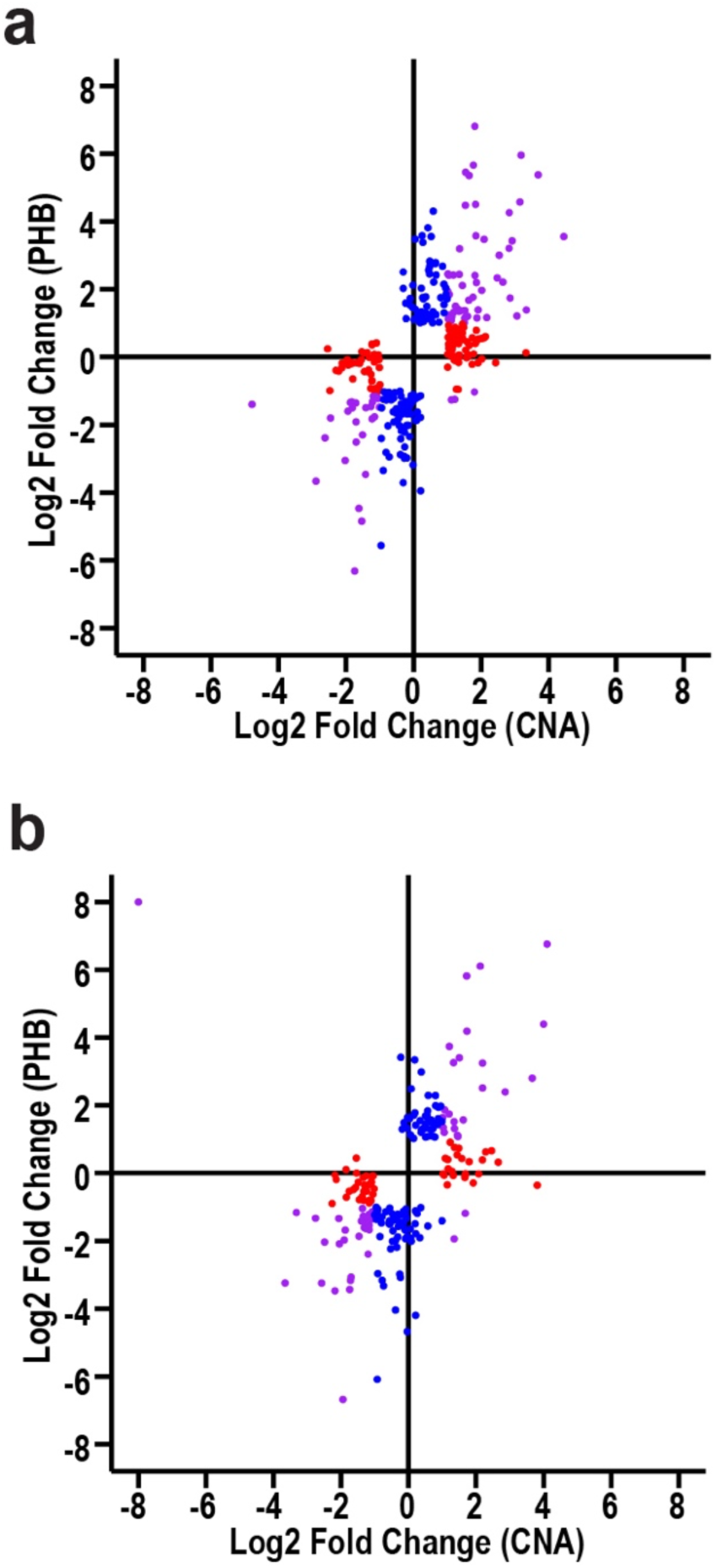
Scatterplots of genes mutually bound by CNA and PHB. **a**, Scatterplot of 317 genes with only mutual affinity binding sites. 81 mutual targets, 107 CNA unique targets, 129 PHB unique targets. **b**, Scatterplot of 208 genes with only mutual binding and CNA higher affinity mutual binding sites. 53 mutual targets, 54 CNA unique targets, 101 PHB unique targets.

**Extended Data Figure 5.**
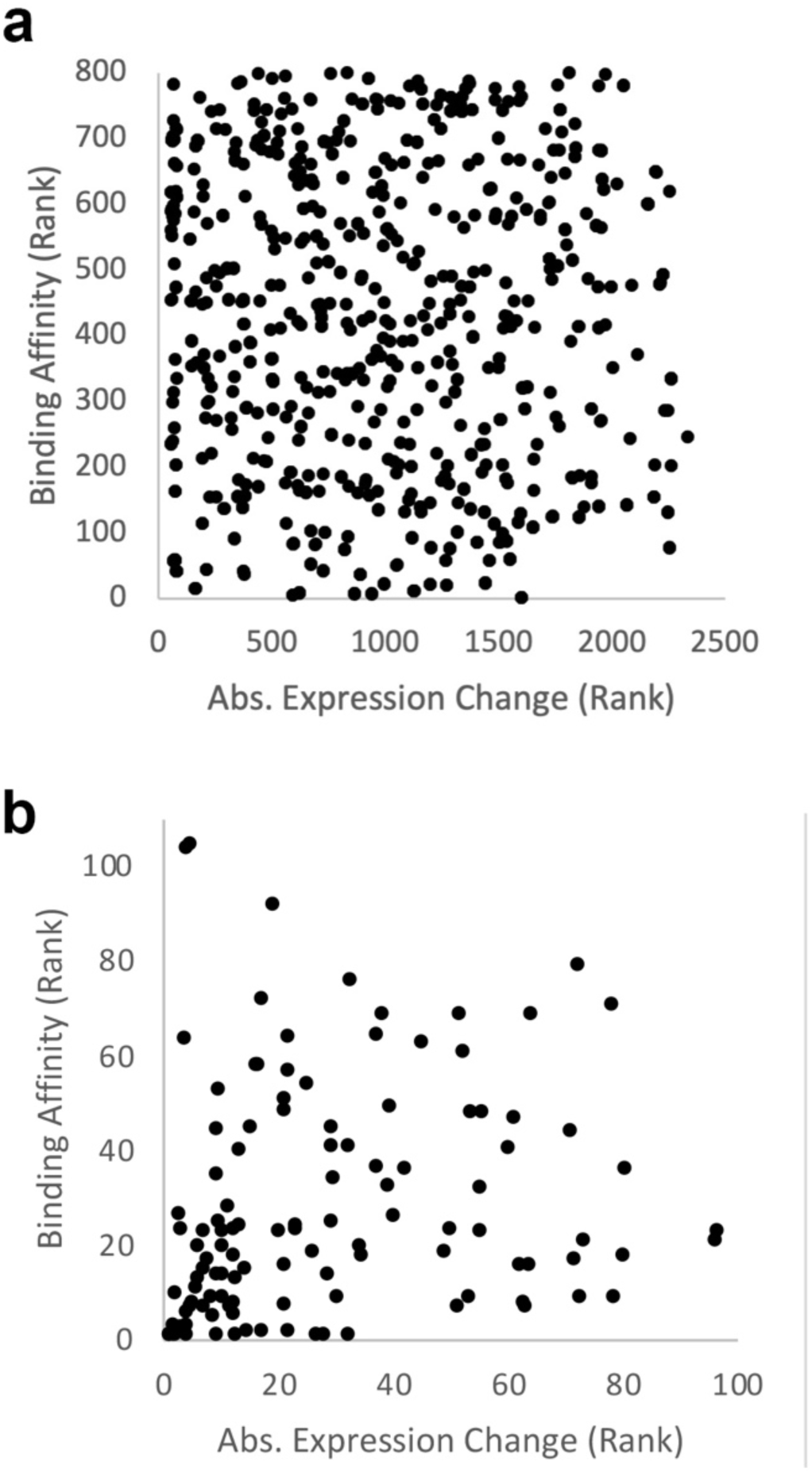
Plots of binding affinity and absolute expression change rankings used in Spearman correlation calculations. **a**, rankings of genes with a singular mutual, CNA higher affinity binding site. Spearman correlation, n = 1,746. R^2^ = .26, p-value = 1. **b**, rankings of genes with a singular mutual, PHB higher affinity binding site. Spearman correlation, n = 119. R^2^ = .26, p-value = 0.99.

**Extended Data Figure 6.**
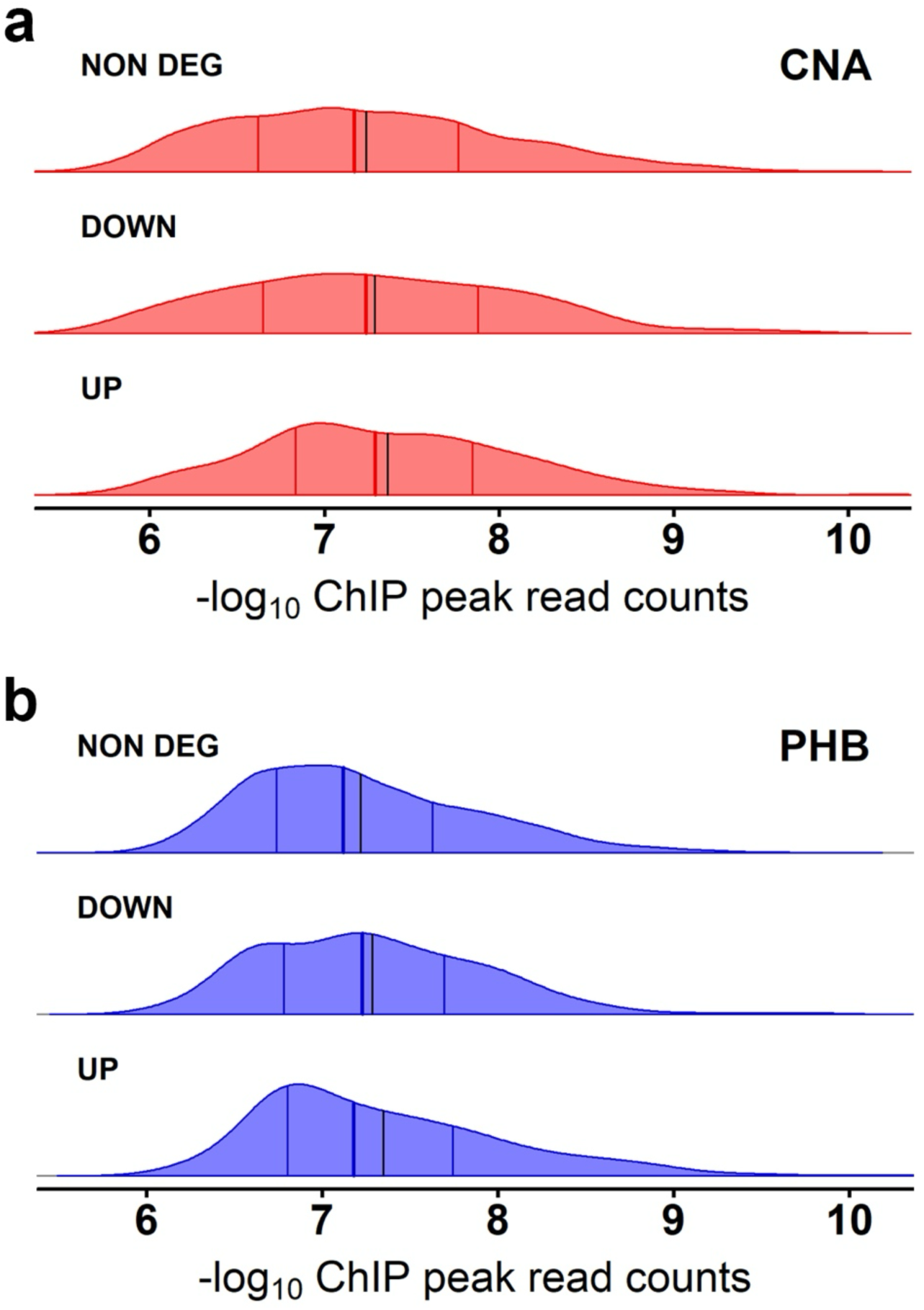
Empirical distribution of ChIP signal at responsive versus non-responsive binding sites. **a**, Ridgeline plots showing distribution of ChIP signal at responsive versus non-responsive sites for CNA. Kruskal-Wallis p-value = 0.006. Mann-Whitney q-values of means of: upregulated versus downregulated genes (0.24); upregulated versus non-regulated genes (0.007); downregulated versus non-regulated genes (0.24). **b**, Ridgeline plots showing distribution of ChIP signal at responsive versus non-responsive sites for PHB. Kruskal-Wallis p-value = 0.007. Mann-Whitney q-values of means of: upregulated versus downregulated genes (0.523); upregulated versus non-regulated genes (0.031); downregulated versus non-regulated genes (0.067). Median values are dark red (CNA) or dark blue (PHB) lines while mean values are black lines in both.

**Extended Data Figure 7.**
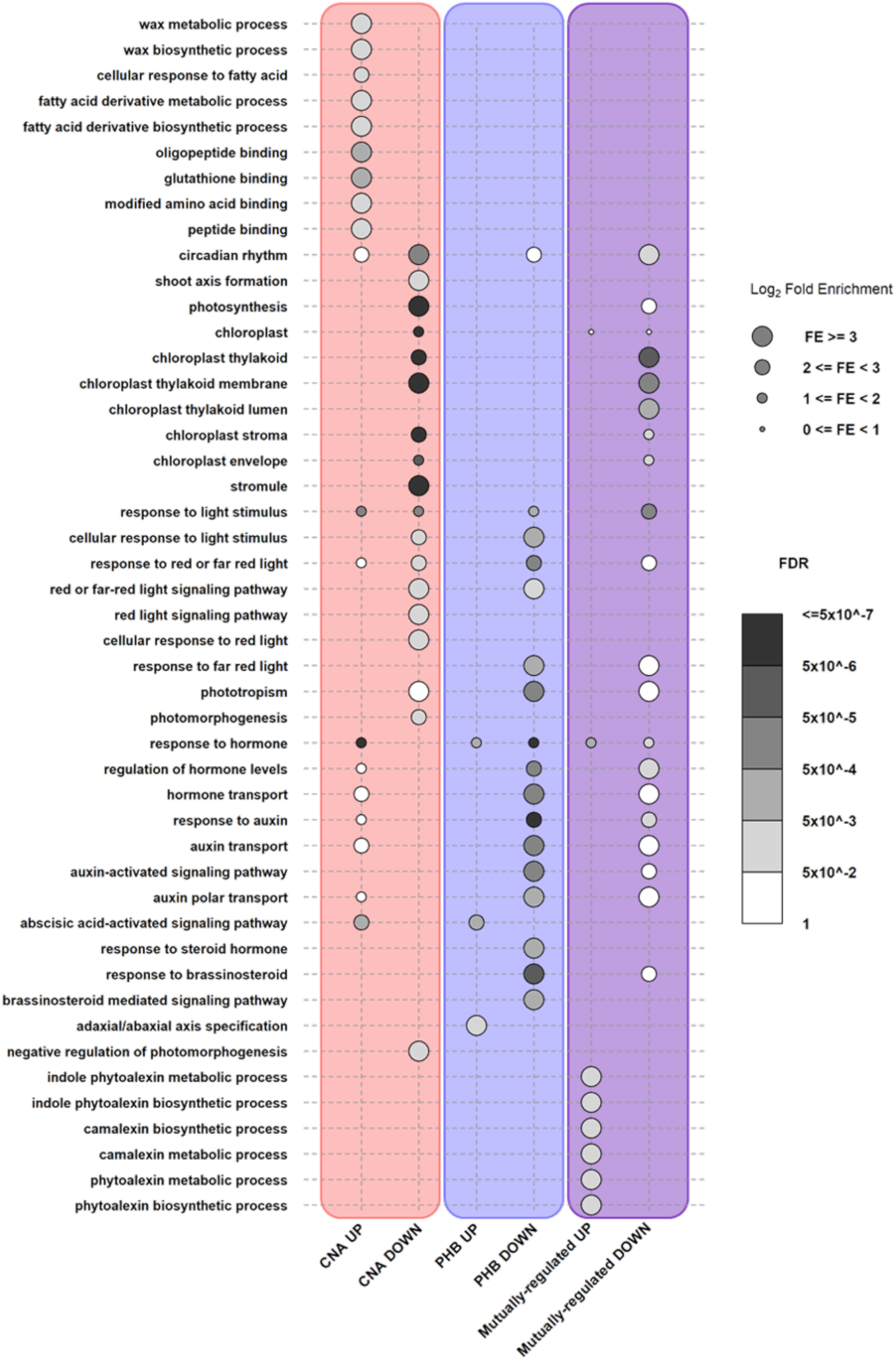
Top Enriched GO Terms. Enriched GO terms for CNA-specific (red), PHB-specific (blue), or mutually regulated (purple) direct targets. The complete output of gene ontology terms can be found in **Supp. Table 6**.

**Extended Data Figure 8.**
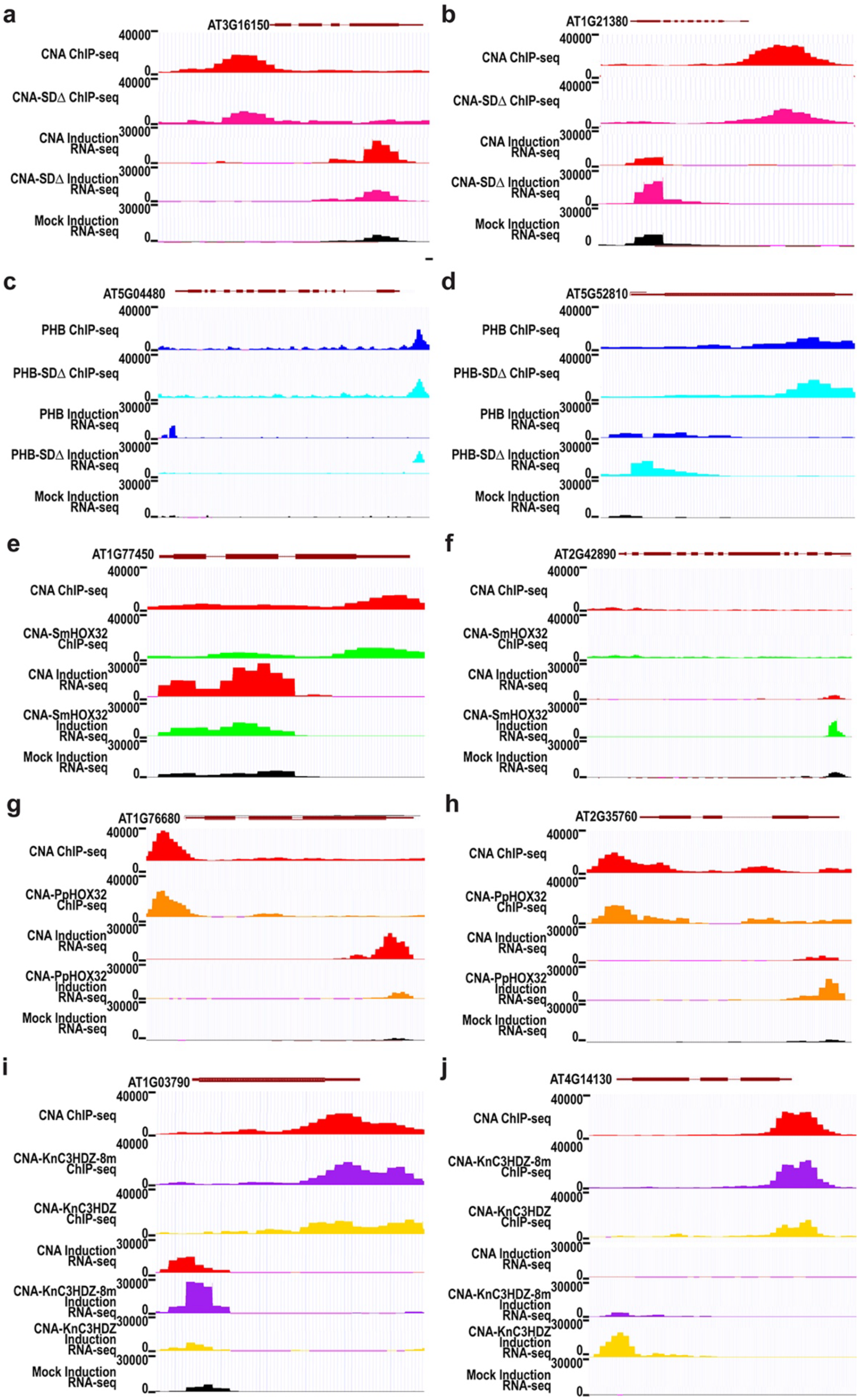
Representative genome browser shots illustrating differential usage of shared binding sites. Genome tracks are presented for the following comparisons: **a, b,** CNA vs CNA-SDΔ; **c, d,** PHB vs PHB-SDΔ; **e, f,** CNA vs CNA-SmHOX32; **g, h,** CNA vs CNA-PpHOX32; **i, j,** CNA vs CNA-KnC3HDZ-8m vs CNA-KnC3HDX.

**Extended Data Figure 9.**
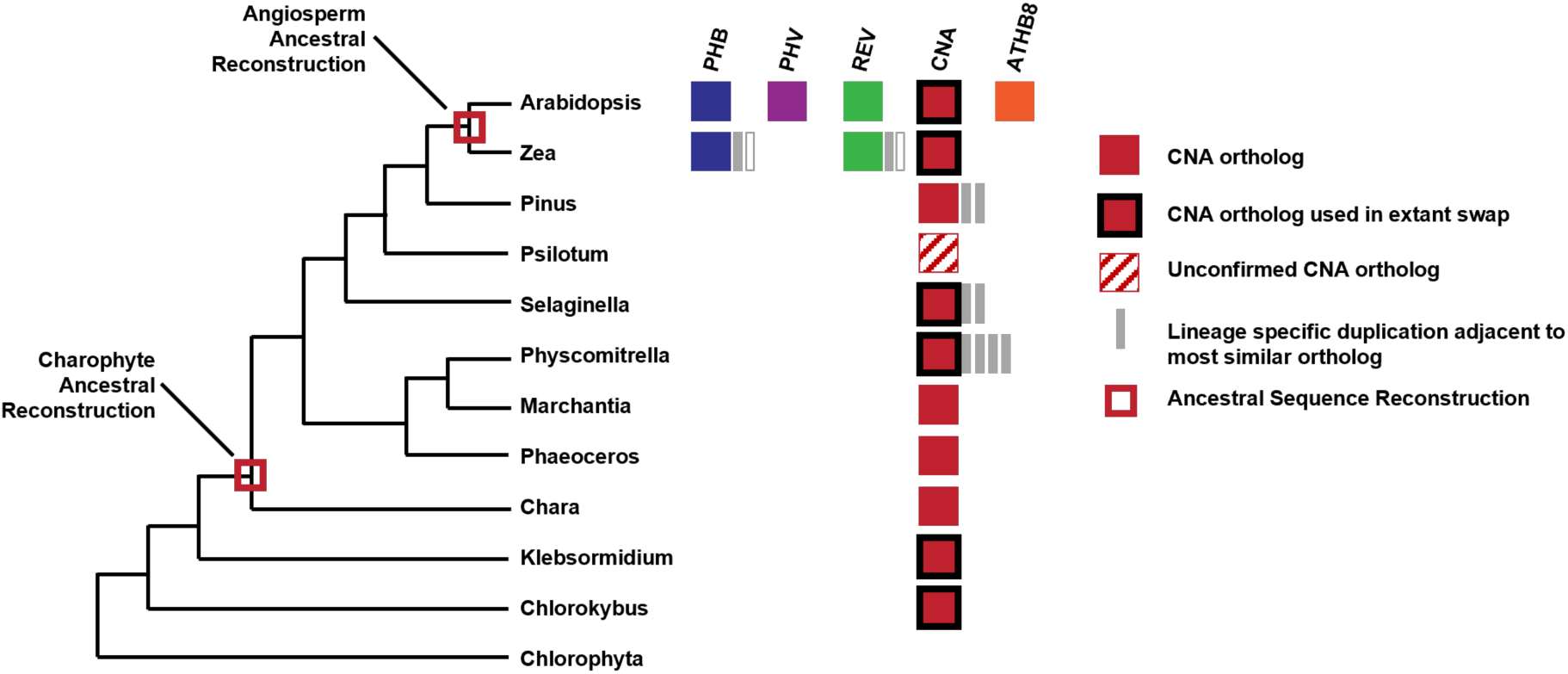
Reconstructed evolutionary history of the HD-ZIPIII family. Duplication and divergence of HD-ZIPIII TFs in multicellular plant species from a CORONA-like ancestral HD-ZIPIII protein.

**Extended Data Figure 10.**
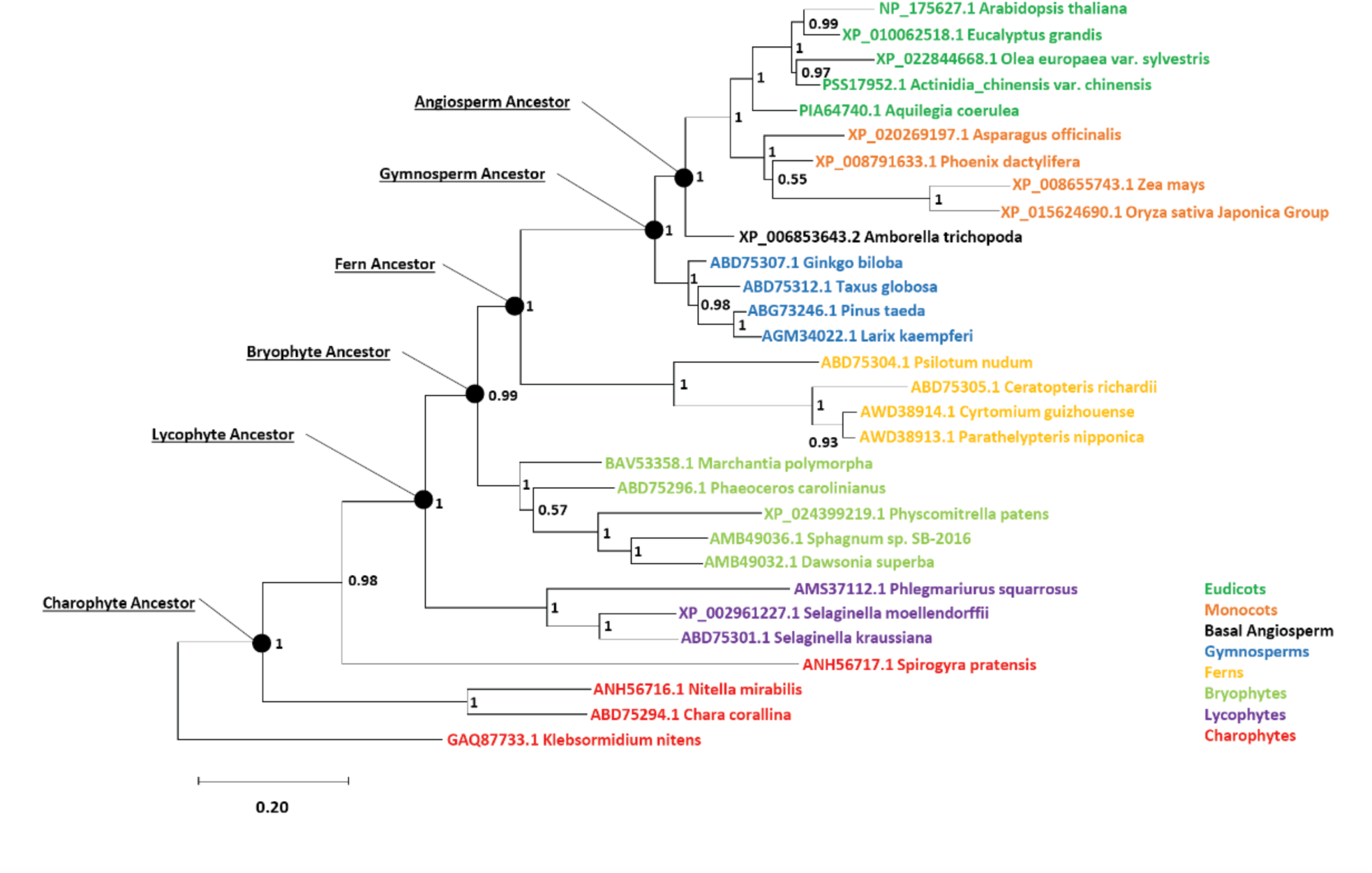
Bayesian inference tree of CORONA orthologs used in ancestral sequence reconstruction.

**Extended Data Figure 11.**
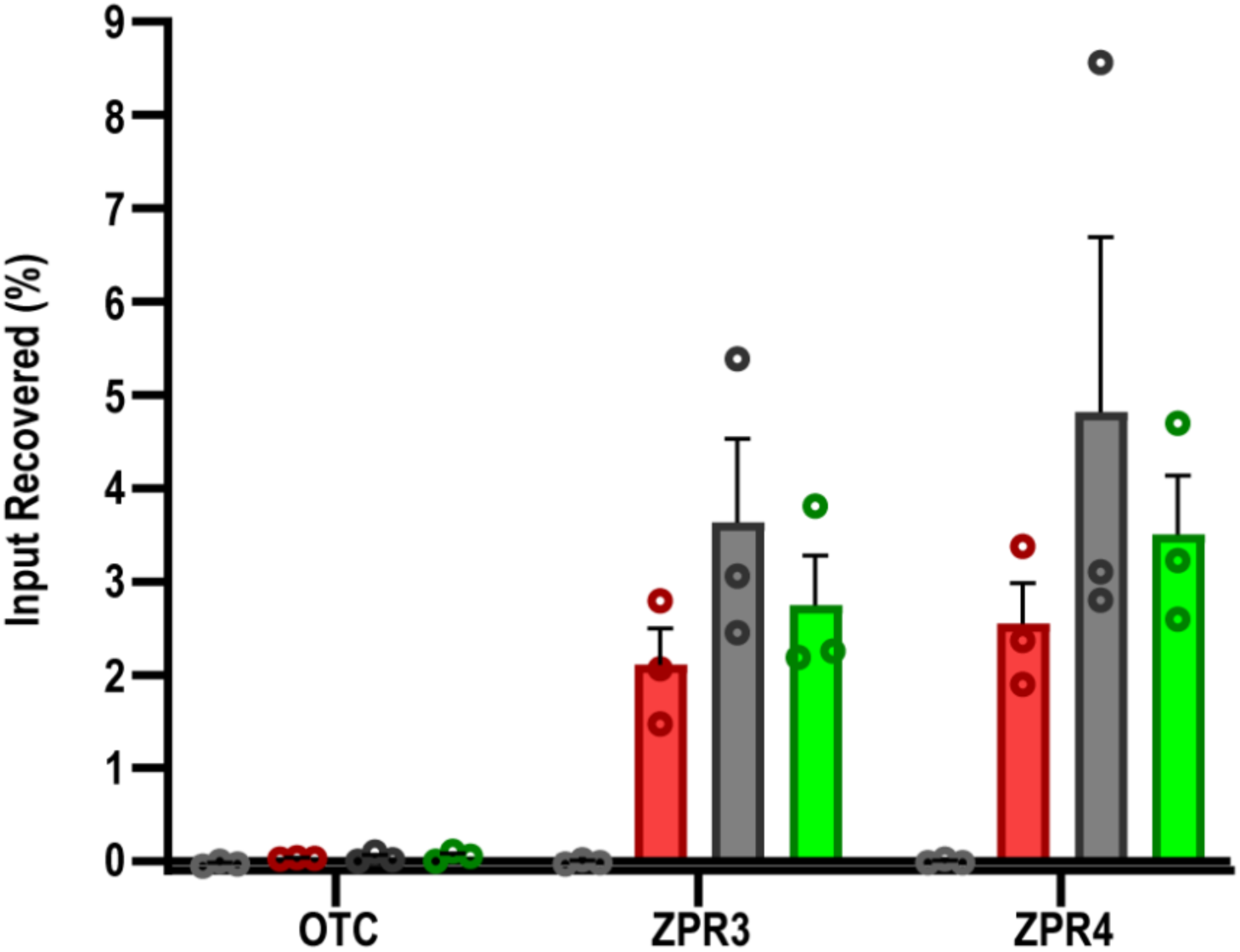
ChIP-qPCR of ZPR3 and ZPR4 CNA binding sites with CNA-CaC3HDZ and CNA-SmHOX32. Both variants show binding despite failure to activate *ZPR3* or *ZPR4*. Light gray = non-transgenic Control, Red= CNA, Dark Gray = CNA-CaC3HDZ, Green = CNA-SmHOX32

**Extended Data Figure 12.**
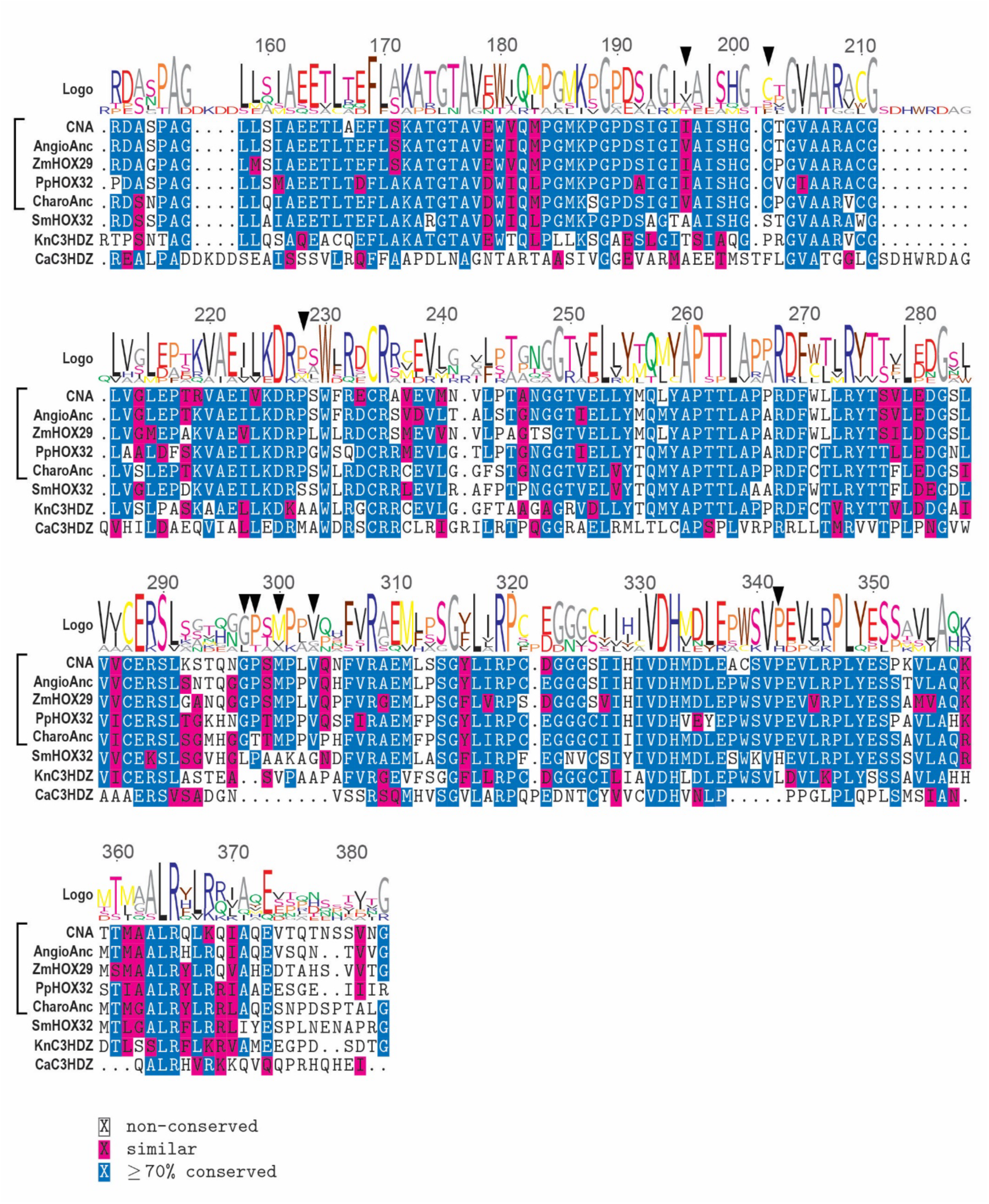
Multiple sequence alignment of START domains used in the chimeric domain swap experiments. Bracketed orthologs are those which condition the pREV:CNA* phenotype in gain of function assays. Blue highlighted residues are those conserved in 5 of the 8 aligned sequences. Residues highlighted in magenta have conserved physiochemical properties. Unhighlighted residues are non-conserved. A triangle over the logo identifies residues which were mutated in the KnC3HDZ START domain to produce CNA-KnC3HDZ_8m. Blue = identical conserved. Magenta = conserved physiochemical. White = not conserved

**Extended Data Figure 13.**
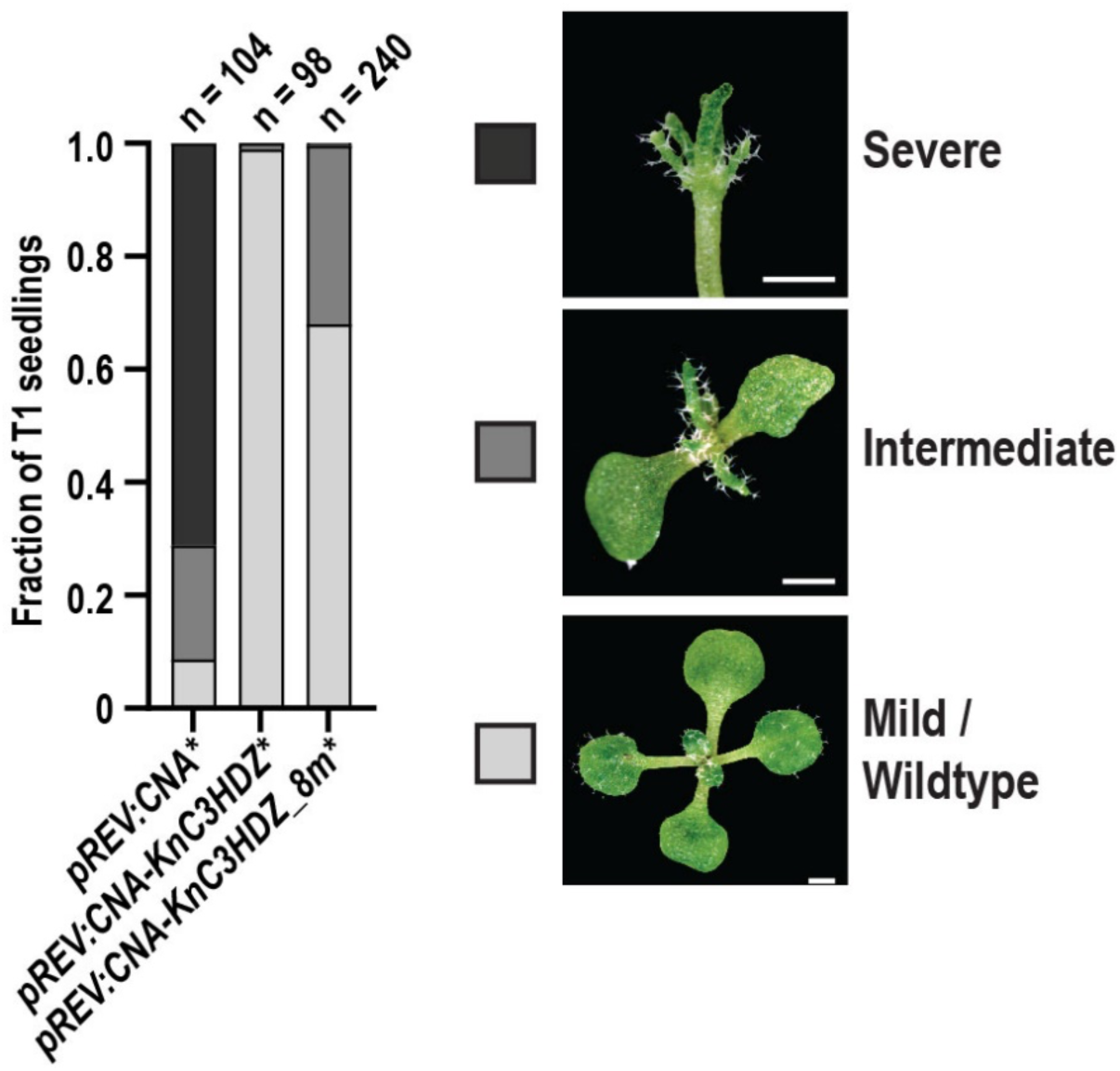
Gain of function assay of T1 transformants of pREV:CNA-KnC3HDZ_8m*. Note images and pREV:CNA* data are replotted/reused from Fig 2b.

